# Disruption of *NAP1* genes supresses the *fas1* mutant phenotype and enhances genome stability

**DOI:** 10.1101/2020.01.03.894170

**Authors:** Karolína Kolářová, Martina Nešpor Dadejová, Tomáš Loja, Eva Sýkorová, Martina Dvořáčková

**Author notes:** these authors contributed equally to the work (Karolína Kolářová, Martina Nešpor Dadejová). Material distribution. The authors responsible for distribution of materials integral to the findings presented in this article in accordance with the policy described in the Instructions for Authors (www.plantcell.org) are: Dr. Martina Dvořáčková; Dr. Martina Nešpor Dadejová.

## Abstract

Histone chaperones mediate assembly and disassembly of nucleosomes and participate in essentially all DNA-dependent cellular processes. In *Arabidopsis thaliana,* loss-of-functions of FAS1 or FAS2 subunits of the H3-H4 histone chaperone complex CHROMATIN ASSEMBLY FACTOR 1(CAF-1) has a dramatic effect on plant morphology, growth and overall fitness. Altered chromatin compaction, systematic loss of repetitive elements or increased DNA damage clearly demonstrate the severity of CAF-1 dysfunction. How histone chaperone molecular networks change without a functional CAF-1 remains elusive. Here we present an intriguing observation that disruption of the H2A-H2B histone chaperone NUCLEOSOME ASSEMBLY PROTEIN 1 (NAP1) supresses *FAS1* loss-of function. The quadruple mutant *fas1nap1;1-3* shows wild-type growth and decreased sensitivity to genotoxic stress. Chromatin of *fas1nap1;1-3* plants is less accessible to micrococcal nuclease and progressive loss of telomeres and 45S rDNA is supressed. Interestingly, the strong genetic interaction between *FAS1* and *NAP1* does not occur via direct protein-protein interaction. We propose that NAP1;1-3 play an essential role in nucleosome assembly in *fas1,* thus their disruption abolishes *fas1* defects. Our data altogether reveal a novel function of NAP1 proteins, unmasked by CAF-1 dysfunction. It emphasizes the importance of a balanced composition of chromatin and shed light on the histone chaperone molecular network.

## INTRODUCTION

Histone chaperones represent an important group of chromatin remodellers, with the ability to directly bind histones and mediate their assembly into nucleosomes (Nakagawa et al., 2001; Takeuchi et al., 2003). This process begins with the interaction of the H3-H4 tetramer with DNA (∼147 bp), followed by addition of H2A-H2B dimers, which completes the formation of the canonical nucleosome. The histone chaperones are also needed for nucleosome disassembly, because many cellular processes (e.g. replication, transcription, DNA repair or recombination) require transient eviction of histones in order to enable easy access of enzymatic machinery to the DNA (Schwabish and Struhl, 2006; Luebben et al., 2010; Zhang et al., 2015). Coordinated interplay between histone eviction, re-loading and *de novo* nucleosome assembly is required for the maintenance of genome stability, in which histone chaperones play an essential role (Lankenau et al., 2003; Mozgova et al., 2010; Zhu et al., 2011; Dewari and Bhargava, 2014). In *Arabidopsis thaliana*, dysfunctions of CHROMATIN ASSEMBLY FACTOR 1 (CAF-1), ANTI-SILENCING FUNCTION 1 (ASF1) or FACILITATING CHROMATIN TRANSCRIPTION (FACT) lead to severe phenotypes, while disruption of HISTONE REGULATOR A (HIRA) or NUCLEOSOME ASSEMBLY PROTEIN 1 (NAP1) are much better tolerated (Exner et al., 2006; Kirik et al., 2006; Liu et al., 2009; Zhu et al., 2011; Duc et al., 2017; Pfab et al., 2018). This suggests that functions of some histone chaperones can be replaced, and that their mutual interactions represent a survival­determining factor. We focused here on the molecular relationship between CAF-1 and NAP1 histone chaperones, which has not yet been investigated.

CAF-1 is a well conserved heterotrimeric complex bringing histones H3.1 and H4 to the replicating DNA, thus contributing to the maintenance of genome stability (Kaufman et al., 1995). Apart from replication-dependent functions, CAF-1 mediates controlled chromatin formation during DNA repair and interacts with several factors involved in homologous recombination (HR), non-homologous end joining or meiosis (Tyler et al., 2001; Endo et al., 2006; Ishii et al., 2008; Almouzni, 2009; Jacob et al., 2014; Muchova et al., 2015; Huang et al., 2018). *Arabidopsis* CAF-1 consists of three subunits called FASCIATA1 (FAS1), FASCIATA2 (FAS2) and MULTICOPY SUPPRESSOR OF IRA1 (MSI1) (Kaya et al., 2001) and mutants of individual subunits have been extensively explored, revealing that a deficiency of *MSI1* is lethal (Bouveret et al., 2006), whereas *fas1* and *fas2* mutant plants survive for nine generations (Mozgova et al., 2010; Pontvianne et al., 2013). The typical *fas1* mutant phenotype is pleiotropic and the most obvious defects include fasciated stems, dentate leaves, short roots and small siliques with a decreased number of seeds connected with fertility problems (Kaya et al., 2001; Exner et al., 2006; Mozgova et al., 2010; Varas et al., 2015). *FAS1* and *FAS2* are cell-cycle regulated, and their deficiency leads to the transcriptional deregulation of several groups of genes, among which histones, S phase-related genes (e.g. B type cyclins, E2F factors), genes involved in DNA repair (*BRCA1, PARP1, Rad51, Rad54*) and plant defence have been of great interest (Bouveret et al., 2006; Exner et al., 2006; Ramirez-Parra and Gutierrez, 2007; Hisanaga et al., 2013; Mozgova et al., 2015). *fas1/2* plants are sensitive to genotoxic stress (methyl methanesulfonate or gamma irradiation), contain higher numbers of DNA damage sites and increased homologous recombination, (Endo et al., 2006; Kirik et al., 2006; Muchova et al., 2015). DNA lesions activate ATAXIA TELANGIECTASIA MUTATED (ATM) target genes and induce G2/M arrest, associated with an early switch to the endocycle and size expansion (Endo et al., 2006; Exner et al., 2006; Ramirez-Parra and Gutierrez, 2007; Hisanaga et al., 2013). All these subcellular changes are reflected in overall fitness and growth abilities of *fas1* and *fas2* mutants, leading to the disorganisation of root and shoot apical meristems (Kaya et al., 2001), abnormal development of the stem cell niche (SCN) (Huang et al., 2018) and a short root phenotype (Muchova et al., 2015; Mozgova et al., 2018). The *FAS1/2* mutation also affects nuclear structure: DAPI stained chromocenters appear smaller and accumulate more of the H3.3 histone variant (Bouveret et al., 2006; Endo et al., 2006; Kirik et al., 2006; Otero et al., 2016). Telomeres and ribosomal rRNA genes (45S rDNA) are systematically lost and silencing of transcriptionally silent information (TSI) loci is released (Mozgova et al., 2010; Duc et al., 2017). Because of these well described phenotypic traits, *fas1/2* mutants represent a valuable experimental model. Considering CAF-1 as a major H3.1 replicative chaperone, the question of how plants cope with its deficiency remains elusive. The most recent studies showed that ALPHA THALASSEMIA-MENTAL RETARDATION X-LINKED (ATRX) and HIRA, chaperones of histone H3.3, are involved in complementary pathways and are able to partially substitute for CAF-1 function, thus maintaining the plant’s viability (Duc et al., 2015; Duc et al., 2017). In accord with this hypothesis, combined mutants (*atrxfas2, fas2hira*) show an aggravated phenotype in comparison to *fas2*.

In this article we focus on the molecular interplay between *FAS1* and genes from the group of chromatin remodellers called *NAP1*. NAP1 family proteins are conserved histone chaperones with apparent pleiotropic functions, as shown in yeast or mammals. NAP1 promotes DNA synthesis-independent nucleosome assembly and disassembly during transcriptional activation (Andrews et al., 2008; Zhang et al., 2015; Chen et al., 2016; Prasad et al., 2016; Dronamraju et al., 2017): it deposits H2A.Z and mediates its cellular transport (Dronamraju et al., 2017); it acts via elimination of non-nucleosomal interactions between H2A/H2B and DNA (Andrews et al., 2010), by shielding the nucleosome-engaged histones (D’Arcy et al., 2013) and by coordination of H1 binding and eviction (Andrews et al., 2008; D’Arcy et al., 2013; Zhang et al., 2015; Shimada et al., 2019). NAP1 is also phosphorylated by cellular casein kinase (CK2), binds to cohesin, and regulates chromatid resolution during mitosis in the fly (Moshkin et al., 2013). *Oryza sativa* NAP1 proteins are cell cycle regulated as well - they bind CycB1, the marker for G2/M and they are phosphorylated by CK2 (Dong et al., 2005). Null mutations of *NAP1* are lethal in mammals (Rogner et al., 2000) and in *Drosophila* (Lankenau et al., 2003), while T-insertion lines in *Arabidopsis* are fully viable (Liu et al., 2009).

Although NAP1 also binds the (H3-H4)_2_ tetramer in yeast and mammals (Andrews et al., 2008), it is considered to be an H2A-H2B histone chaperone in plants, where binding properties of NAP1 and H3-H4 have not been investigated (Dong et al., 2005).

The *Arabidopsis NAP1* family comprises four *NAP1* genes (*NAP1;1-4*) and two *NAP1-RELATED PROTEIN* genes (*NRP1, NRP2*). NAP1;1-3 proteins are ubiquitously expressed, form homodimers and heterodimers and localize abundantly to the cytoplasm (Liu et al., 2009). *NAP1;4* encodes a shorter protein lacking canonical sequences and is only weakly expressed in roots and pollen grains (Liu et al., 2009; Zhou et al., 2016). Loss-of-function mutations of individual *NAP1* genes do not affect plant growth and development under standard laboratory conditions (Gao et al., 2012). Triple mutants *nap1;1-3* are sensitive to UV-C irradiation and have reduced levels of somatic homologous recombination (Gao et al., 2012).

*NRP*s are related to the yeast VACUOLAR PROTEIN SORTING (Vps75 and human SET/TAF proteins and participate in nucleosome assembly and histone binding (Zhu et al., 2006; Tang et al., 2008; Bowman et al., 2011; Bowman et al., 2014; Kumar et al., 2019). *NAP1* and *NRPs* act in a synergetic molecular pathway to regulate HR, however, there is a lack of information on which step/s of it they participate in (Zhu et al., 2006; Zhou et al., 2016). The multiple mutants of *nap1;1-4nrp1-2* are largely sensitive to genotoxic stress, but are otherwise viable (Zhou et al., 2016). Unlike NAP1, which is considered to be an H2A/HB histone chaperone shuttling between the cytoplasm and nucleus, NRP proteins are nuclear factors forming homo-and heterodimers and binding all histones *in vitro* (Liu et al., 2014; Gonzalez-Arzola et al., 2017).

Genetic interaction was previously described for *FAS2* and *NRP1-2* genes in HR and maintenance of the stem cell niche (Gao et al., 2012; Huang et al., 2018). The hyper-recombinogenic phenotype of *fas2* plants contrasts with *nrp1-2* and *nap1;1-3* mutants. In the triple mutant *fas2nrp1-2,* somatic HR shows similar levels to that of the double mutant *nrp1-2,* indicating that the *nrp1-2* mutation overrides the CAF-1 dysfunctional effect.

Here we investigated the relationship between FAS1 and NAP1 histone chaperones using a genetic approach and created a *fas1nap1;1-3* quadruple mutant line. We show that *fas1nap1;1-3* lacks most of the phenotypic abnormalities found in *fas1* single mutants, such as worsened plant fitness and growth. We also analysed the influence of *nap1;1-3* mutations on the overall chromatin state, 45S rDNA and telomere maintenance, and the DNA damage response. With respect to our findings, we propose a direct link between NAP1;1-3 and the FAS1 pathway, where NAP1 proteins are epistatic over FAS1 and contribute to the mutant phenotype in *fas1* lines.

## RESULTS

### NAP1 proteins contribute to penetration of the *fas1* phenotype

To discover a possible relationship between FAS1 and NAP1;1-3 histone chaperones, and elucidate their roles in chromatin maintenance, we crossed the *fas1-4* single mutant (G1) with the *nap1;1nap1;2nap1;3-2* triple mutant (*m123-2*, adopted from (Liu et al., 2009)), thereby creating the *fas1-4nap1;1nap1;2nap1;3* (*fas1m123-2*) quadruple mutant. Parental *A. thaliana* plants carrying a T-DNA insertion in all three *NAP1;1-3* genes *(m123-2*) grow relatively well under standard laboratory conditions without any apparent phenotypic changes (Liu et al., 2009), while dysfunction of the *FAS1* gene causes growth and developmental problems and reduced fertility (Kaya et al., 2001; Kirik et al., 2006; Mozgova et al., 2010; Hisanaga et al., 2013; Varas et al., 2017).

We found that simultaneous loss-of-function of *NAP1;1, NAP1;2, NAP1;3* and *FAS1* genes barely affects plant growth and morphology under standard growth conditions. The *fas1m123-2* mutant plants do not show serrated leaves, as do *fas1* plants, and they resemble wild-type (WT) (Figure 1A). In addition to improved plant growth, *fas1m123-2* plants are fully fertile, in comparison to *fas1*. We performed an extended analysis and investigated the shape of anthers and the number of viable pollen grains using a modified Alexander staining protocol (Peterson et al., 2010). *fas1* plants revealed an altered shape of anthers and reduced number of pollen grains compared to the WT. These developmental changes were not observed in *fas1m123-2* or *m123-2* mutant plants (Figure 1B, C). In addition, no aborted seeds were observed in these lines, suggesting that the fertility problem observed in *fas1* was eliminated in *fas1m123-2*. In conclusion, *fas1m123-2* does not show any growth defects typical for *fas1* mutants (Figure 1A-C).

**Figure 1.**
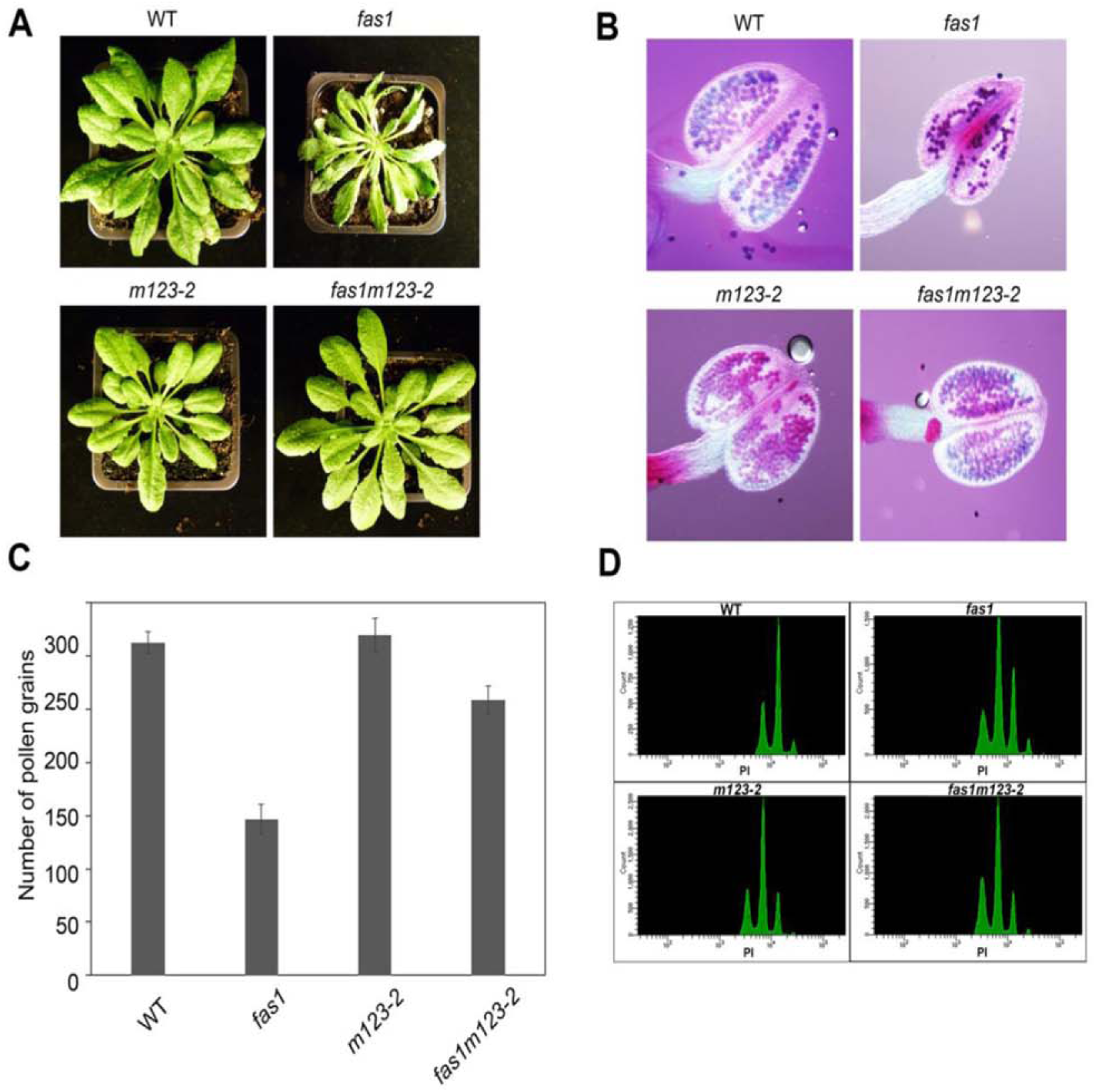
Morphology of *fas1m123-2* plants is more similar to the WT. A) Images of WT, *fas1, m123-2* and *fas1m123-2* plants collected after 3-weeks growth under short-day conditions in controlled growth chambers. Visible morphological changes are apparent only in *fas1* mutant plants but not in *fas1m123-2* plants. B) Pioidy distribution of WT, *fas1, m123-2* and *fas1m123-2* plants measured by flow cytometry in the nuclei of 22-d-old leaves. Higher levels of 8C and 16C nuclei occur in *fas1* but not in *fas1m123-2* plants. C) Alexander staining of pollen grains in WT, *fas1, m123-2* and *faslm123-2* mutant lines. Bright field images of the stained anthers, the violet colour represents viable spores, while green indicates aborted spores. The majority of spores in individual mutants are viable. D) Graphical representation of the average number of pollen grains per anther in WT and mutant lines. Altered shape of anther and lower amount of pollen grains is present in *fas1* mutants. Pollen grains from three anthers from each line were counted.

Plant growth depends on efficient cell cycle progression and requires a good balance between proliferation and cell expansion. Leaves of the *fas1* mutant display a high ploidy phenotype, which is compensated by cell enlargement (so called compensation phenotype) (Endo et al., 2006; Ramirez-Parra and Gutierrez, 2007; Hisanaga et al., 2013). To determine whether the improved plant growth in *fas1m123-2* relates to changes in pioidy distribution we measured the dna content in nuclei isolated from leaves of 22-d-old plants (Figure 1D, Supplemental Figure 1). All lines revealed three major peaks - 2C, 4C and 8C with varied intensities (Figure 1D). In *fas1,* the 8C peak had a higher intensity, as expected (Figure 1D) and 16C and 32C peaks were clearly detected (Supplemental Figure 1). The pattern of ploidy distribution in *fas1m123-2* was more similar to the pattern in *m123-2* plants or WT than to *fas1* (Figure 1D, Supplemental Figure 1) and it showed a lower abundance of higher ploidy (8C to 32C) nuclei.

### Expression levels of *NAP1;1-3* are elevated in *fas1* mutants

We further asked what are the mRNA levels of *NAP1* genes in *fas1,* as *NAP1;1-3* disruption in *fas1m123-2* caused such a remarkable effect. The studies performed on *fas1/2* so far did not find any *NAP1* histone chaperones among the list of deregulated genes (Bouveret et al., 2006; Huang et al., 2018). To our surprise, the RT-qPCR performed on *fas1* seedlings revealed induction of all three *NAP1* genes in *fas1* compared to WT (Figure 2A) - approximately seven-fold higher transcript levels of *NAP1*;*1*, two-fold of *NAP1*;*2* and three-fold of *NAP1*;*3.* Transcript levels of *NAP1;1-3* were negligible in *m123-2*, as expected from disruption of *NAP1* genes in T-DNA insertion lines. In the *fas1m123-2* mutant, abundance of *NAP1;1* and *NAP1;3* transcripts decreased. *NAP1;2* was not detected in leaves of individual *fas1m123-2* plants (Figure 2B), while some *NAP1;2* transcription was detected in seedlings (Figure 2A). Since we genotyped both the *nap1;1-3* parental line obtained from Liu *et al*. (Liu et al., 2009) and the *fas1nap1;1-3* quadruple line as heterozygous for the *NAP1;2* gene, a mixture of seedlings used for a particular qPCR could also segregate to WT plants in respect of the *NAP1;2* locus in *fas1m123-2*, further explaining detectable expression of *NAP1;2* in seedlings.

**Figure 2.**
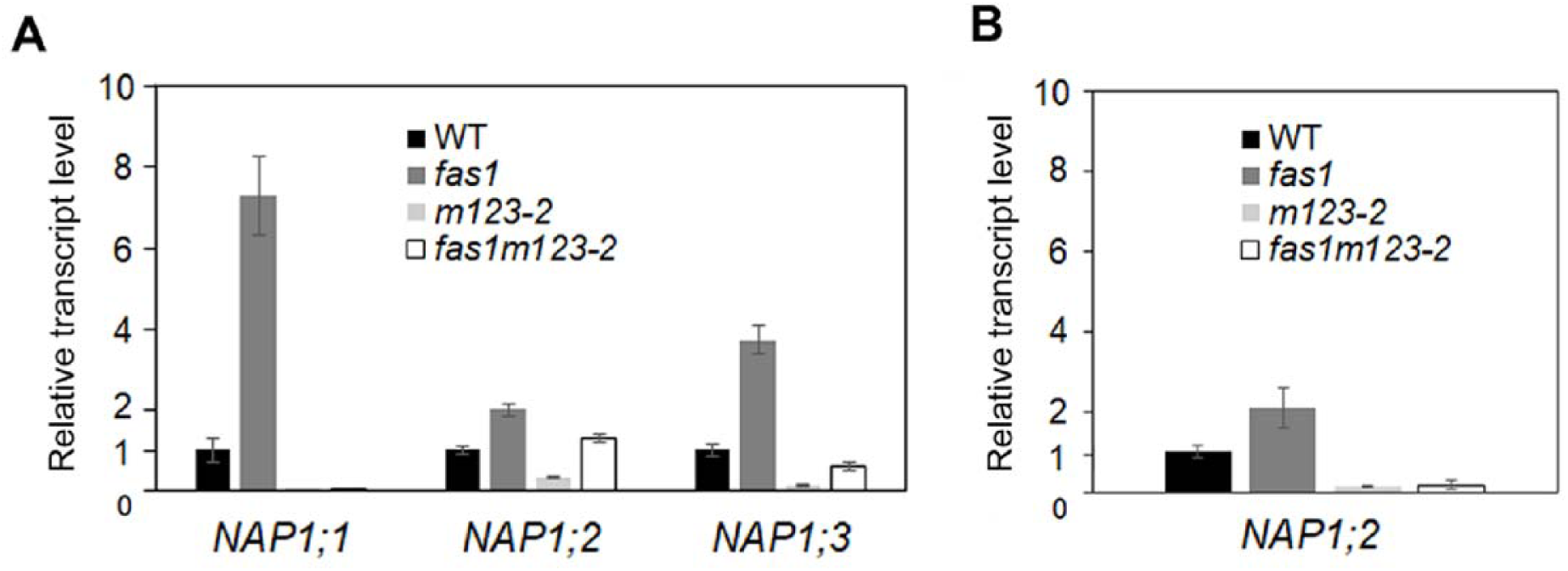
*NAP1* genes are overexpressed in *fas1*. A) Transcript levels of *NAP1;1, NAP1;2* and *NAP1;3* genes were determined in 10-d-old seedlings of WT and mutant lines *(fas1, m123-2* and *fas1m123-2).* The *fas1* mutant shows higher expression of all *NAP1* genes than WT plants. Error bars indicate standard deviations calculated from three biological replicates, ubiquitin 10 was used as a reference gene. B) Relative transcript levels of *NAP1;2* gene were determined in leaves of 4-w-old plants of WT and mutant lines (*fas1, m123-2* and *fas1m123-2).* The *fas1* mutant shows two-fold higher expression of NAP1:2 than WT plants, expression in *m123-2* and *fas1m123-2* mutants is negligible. Error bars indicate standard deviations calculated from three biological replicates, ubiquitin 10 was used as a reference gene.

### The *fas1m123-2* mutant is more tolerant to genotoxic stress than *fas1*

One of the described phenotypes of *fas1* is the increased sensitivity to the DNA damaging chemical, methyl methanesulfonate (MMS) (Muchova et al., 2015). MMS is an alkylating agent that stalls replication forks, blocks DNA polymerases and inhibits DNA synthesis (Slamenova et al., 1990). As the next step, we tested whether *fas1m123-2* plants remained sensitive to MMS. To assess the response to this genotoxin, root lengths of WT, *fas1, m123-2* and *fas1m123-2* were measured in 10-d-old seedlings. In agreement with previous studies (Exner et al., 2006; Huang et al., 2018), *fas1* mutants showed a short root phenotype on ½ MS medium without MMS (Figure 3A) and further growth inhibition on medium supplemented with 125 μΜ MMS (Figure 3B,C). In contrast, *fas1m123-2* seedlings did not show any retardation of root growth on control plates nor after MMS treatment (Figure 3A-C). The root lengths in *fas1m123-2* were similar to that in the *m123-2* mutant line and only slightly shorter than in WT. In conclusion, *fas1m123-2* seedlings were no longer sensitive to MMS, thus the *fas1* phenotype was compensated for in the quadruple mutant.

**Figure 3.**
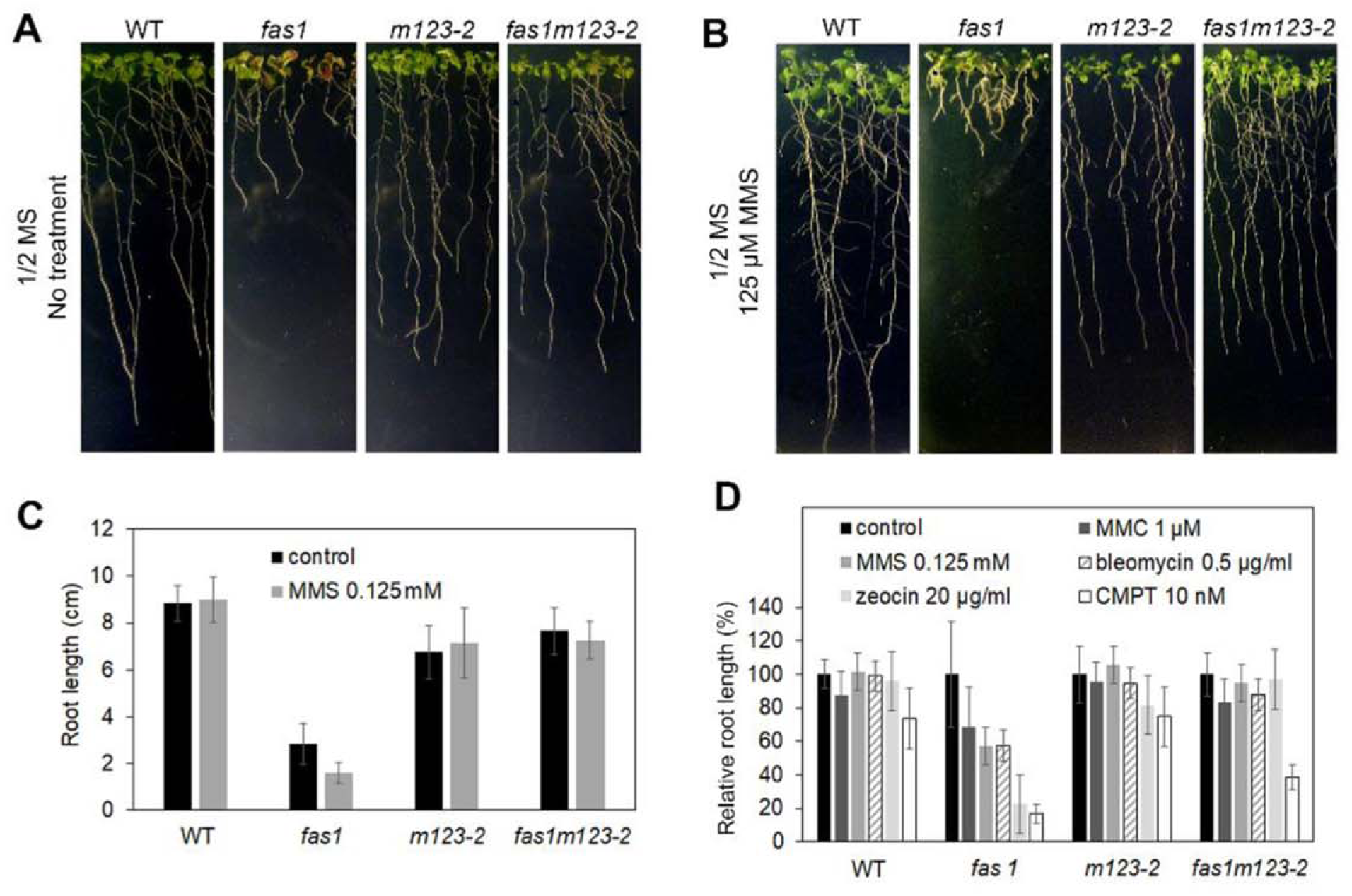
*fas1m123-2* plants are not sensitive to genotoxic agents. A) 10-d-old WT, *fas1, m123-2* and *fas1m123-2* plants grown on plates with ½ MS medium (no treatment). B) 10-d-old WT, *fas1, m123-2 and faslm123-2*plants grown for 3 days on plates with ½ MS medium and for additional 7 days on ½ MS medium containing 125 μΜ MMS. C) Absolute root lengths of WT, *fas1, ml23-2* and *fas1m123-2* plants on control ½ MS medium and on medium supplemented with 125 μΜ MMS; 40 plants of each line were used in this analysis. The results are summarised in the chart, error bars indicate standard deviations. D) Relative change of root lengths of WT, *fas1, m 123-2* and *fas1m123-2* plants grown on control ½ MS medium (100%) and on media supplemented with given concentrations of MMS, MMC, bleomycin, zeocin and camptothecin. Root lengths of 40 plants were measured on seedlings grown for 7 days on genotoxic agents. The results are summarised in the chart, error bars indicate standard deviations.

We next investigated whether *fas1m123-2* acquired a specific resistance to MMS or a more general tolerance to any type of genotoxic stress. Since MMS causes double-strand-breaks (DSB) only after replication (Lundin et al., 2005), we selected DNA damaging agents affecting various steps of DNA metabolism: a) camptothecin (CPT), that binds and stabilises the topoisomerase I (TOP1) - DNA cleavage complex, consequently leading to a collision with the replication fork in actively dividing cells and DSB (Pizzolato and Saltz, 2003); b) bleomycin and zeocin, two commonly used radiomimetic agents inducing DSB, chromosomal aberrations or translocations (Pecinka and Liu, 2014); and c) mitomycin C (MMC), a bifunctional alkylating agent able to form inter-strand crosslinks and therefore inhibit replication and transcription (Pecinka and Liu, 2014; Manova and Gruszka, 2015).

The concentrations of all inhibitors were adjusted to levels that did not affect WT plants. Due to the *fas1* short root phenotype, it is difficult to evaluate changes in the absolute root lengths and to compare growth in the presence of different inhibitors (Supplemental Figure 2). We thus decided to calculate relative changes of root length in WT and all mutant lines, where the root length on the control plate was always set as 100%. We found that *fas1* mutants were equally sensitive to MMS, MMC and bleomycin, showing about 50% retardation of root growth (Figure 3D). Zeocin and CPT had even stronger effects, causing 80% retardation of root growth. In comparison to *fas1, fas1m123-2* mutants acquired resistance to MMS, MMC, bleomycin and zeocin, indicating that the disruption of *NAP1* genes in *fas1* causes general tolerance to genotoxic stress. An interesting effect was observed on medium containing CPT; *fas1m123-2* plants grown on CPT were not as sensitive as *fas1,* but did not reach full CPT tolerance, as observed in WT or *m123-2* (Figure 3D).

### *fas1m123-2* shows lower levels of programmed cell death in root tips

The CAF-1 complex was shown to regulate cell type-specific gene expression in the root apical meristem (Kaya et al., 2001) and to prevent programmed cell death in the stem cell niche (Huang et al., 2018). We show here that retardation of root growth that occurred in *fas1* was abolished in *fas1m123-2* plants and sensitivity to genotoxic stress was decreased. We further asked whether the formation of apoptotic lesions in *fas1* mutants (Kaya et al., 2001) depended on NAP1 proteins, and thus carried out a propidium iodide (PI) analysis (Fulcher and Sablowski, 2009). Propidium iodide lightly stains the cell walls of viable cells, while it intensively labels the cellular interior when the integrity of the cell membrane is impaired. Whole seedlings were used for PI staining and the apical root meristem was imaged using confocal microscopy. We observed that roots of untreated WT and *m123-2* plants occasionally contained small PI-stained areas, while more than 50% of *fas1* roots showed large apoptotic lesions, as expected (Figure 4A and 4B). *fas1m123-2,* in contrast, showed small lesions in ∼25% of roots - the pattern was more similar to WT than to *fas1* plants. This improvement in root growth in *fas1m123-2* was correlated with the lower number of damaged cells in the root tips. We then investigated the frequency of apoptotic lesions after treatment with CPT. The resistance of *fas1m123-2* to CPT (Figure 3D) was not complete, suggesting that CPT would be a good choice for tracking even small changes. We tested acute and chronic responses to CPT. To assess the acute response, *Arabidopsis* 4-d-old seedlings were treated with CPT in liquid medium for 24 h (Figure 4A), while the chronic response was determined using plants grown on agar plates containing CPT for 6 days (Figure 4B). This experiment revealed clear differences between individual lines. Acute CPT treatment induced a certain level of apoptotic lesions in all four genotypes tested. In WT and *m123-2,* relatively small increases in PI lesions were observed compared to untreated plants. The highest response to CPT was detected in *fas1m123-2 —* 50 % of the roots formed PI lesions (∼20% small, 30% large), but the overall highest level of damage was observed in *fas1*, where all roots examined showed small or large lesions (Figure 4A). Chronic CPT treatment resulted in a different pattern (Figure 4B). In *fas1,* root growth and normal development were completely inhibited compared to *fas1m123-2*, where they remained unchanged (Figure 4B). Although the proportion of PI stained roots increased in *fas1m123-2,* the observed ratio became more similar to the *m123-2* mutant (40% small, 60% large lesions) than to *fas1* or WT. In WT, 50% of roots remained intact even after 6 days of CPT treatment, indicating a relatively mild effect. We concluded that NAP1 proteins participated in the chronic response to genotoxic stress and their disruption in *m123-2* mutants caused increased cell death in the root SCN, in comparison to WT. NAP1 dysfunction in *fas1* (*fas1m123-2*), on the other hand, largely improved tolerance to CPT, indicating that both histone chaperones contributed to CPT-induced DNA repair and their imbalance in individual mutants (*fas1* or *m123-2*) was much more deleterious than *fas1m123-2*.

**Figure 4.**
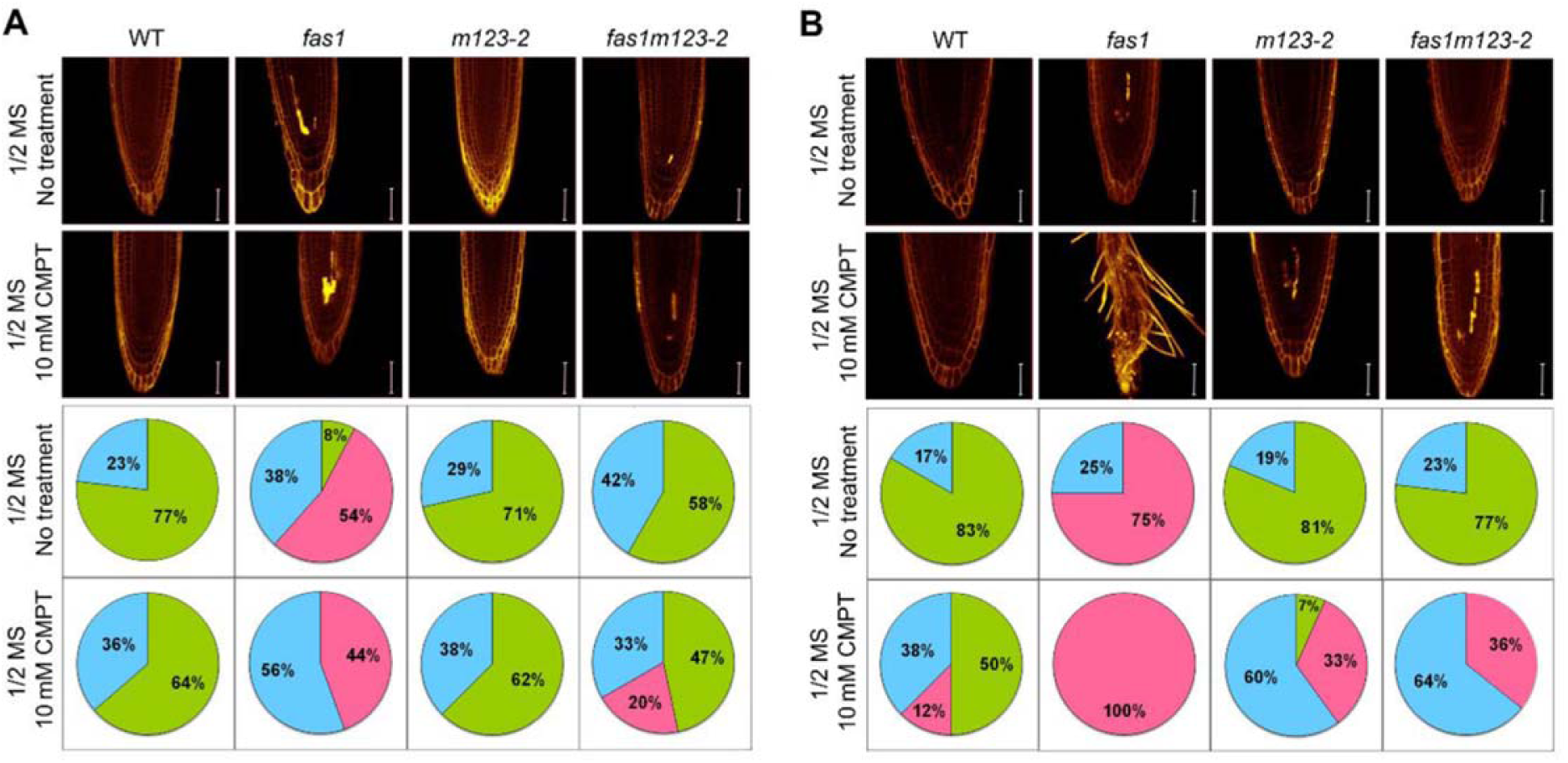
NAP1 proteins participate in the regulation of the root stem cell niche. A) Propidium iodide staining of the apical root meristem in WT, *fas1, ml23-2* and *fas1m123-2*seedlings after 24 h treatment with/without 10 nM camptothecin (CPT) (top panels) and graphical representation of the occurrence and extent of cell death (bottom panels). 20 root tips of each line were examined: green = no damage, blue = small damage, magenta = large damaged areas. Scale bars: 200 pm. B) Propidium iodide staining of the apical root meristem in WT, *fas1, ml23-2* and *fas1m123-2* seedlings after 6 days of treatment with/without 10 nM CPT (top panels) and graphical representation of the occurrence and extent of cell death (bottom panels). 20 root tips of each line were examined: green = no damage, blue = small damage, magenta = large damaged areas. Roots of *fas1* mutants were destroyed after 6 days of treatment with CPT. Scale bars: 200 μm.

### Expression of *BRCA1* and *PARP1* is decreased in *fas1m123-2* compared to *fas1* mutants

Several independent studies pointed out that *fas1* and *fas2* mutants show increased levels of expression of *BRCA1*, *PARP1* and *Rad51*, three important DNA repair genes (Bouveret et al., 2006; Ramirez-Parra and Gutierrez, 2007; Hisanaga et al., 2013; Huang et al., 2018). Since we observed that the effect of DNA damaging agents was less deleterious in *fas1m123-2* than in *fas1, we* investigated whether this effect was related to changes in *BRCA1* and *PARP1* expression. The expression analysis using RT-qPCR was performed in WT, *fas1, m123-2* and *fas1m123-2* seedlings grown under normal conditions and compared to expression in seedlings treated with CPT (Figure 5). *BRCA1* and *PARP1* showed comparable levels of expression in WT and the *m123-2* mutant, with induction of both genes after CPT treatment. The most profound differences were observed in *fas1* plants, with *BRCA1* and *PARP1* levels being significantly elevated 6 and 4-fold, respectively, when compared to WT. Moreover, *fas1* plants did not show induction of gene expression after CPT treatment. In contrast, in *fas1m123-2, BRCA1* and *PARP1* expression was ca. 3-fold higher than WT - about a half of their expression in *fas1*. Interestingly, transcription of these two genes was not further induced by CPT treatment, similarly to *fas1* (Figure 5). Our data demonstrate that constitutive activation of DNA repair occurred in both mutants studied and that the plateau of *PARP1* and *BRCA1* induction was lower in *fas1m123-2*.

**Figure 5.**
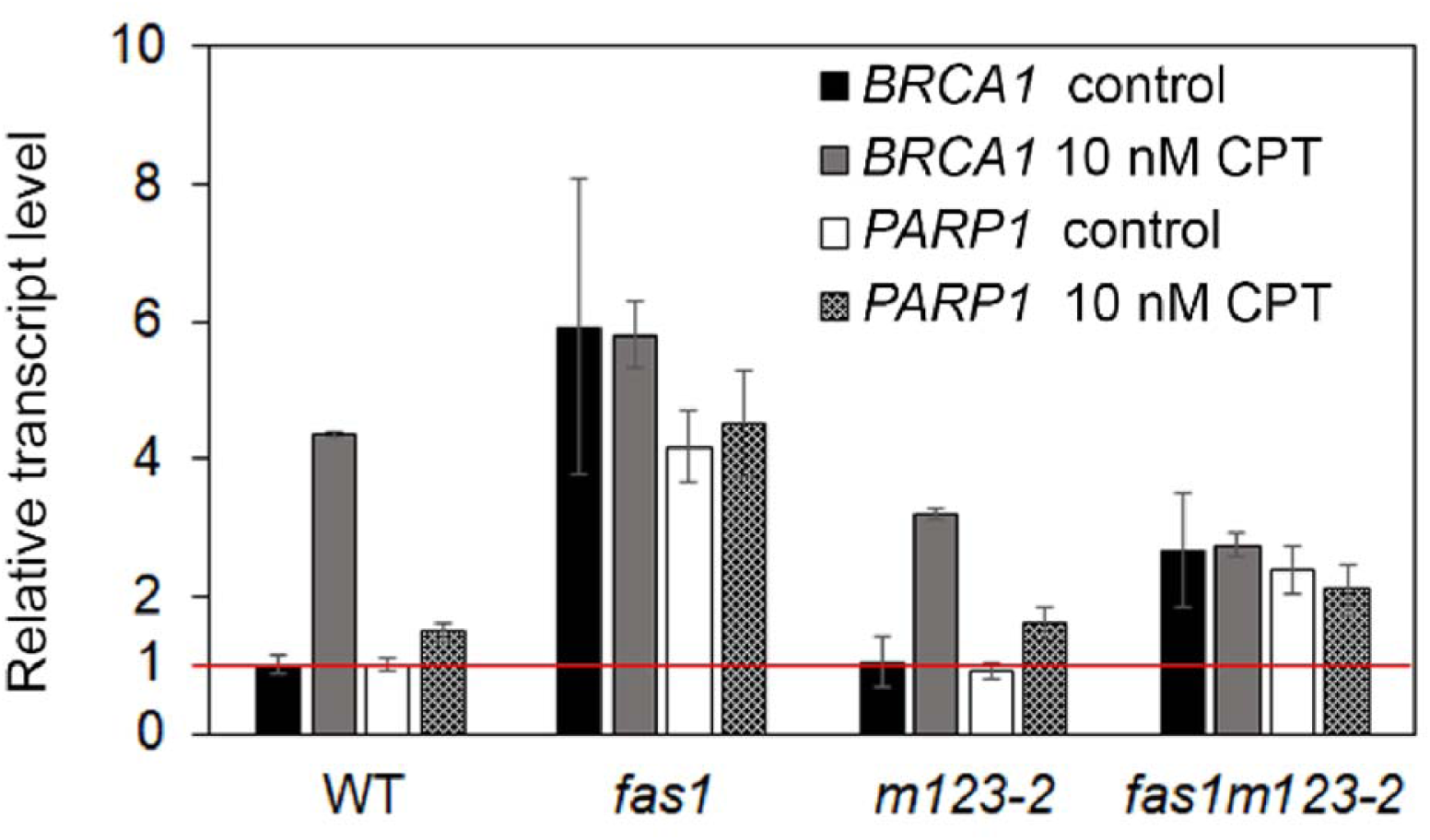
*BRCA1* and *PARP1* mRNA levels are altered *fas1m123-2.* Transcript levels of DNA damage response genes *BRCA1* and *PARP1* in 10-d-old seedlings of WT, *fasl, m123-2* and *fas1m123-2* on ½ MS medium (control) and on medium supplemented with 10 nM CPT. Relative expression levels with WT set up as 100% −1 and using ubiquitin 10 as a reference are shown. Three biological replicates were analysed, three technical replicates each, error bars indicate standard deviations.

### DNA damage signalling occurs normally in *fas1m123-2*

Considering the lower sensitivity of *fas1m123-2* to genotoxic agents and decreased levels of DNA damage-response genes, *BRCA1* and *PARP1,* we wondered whether DNA damage signalling would be induced with the same efficiency in the quadruple mutant. We employed an established marker of DNA damage signalling, γ-H2A.X (Friesner et al., 2005; Zhou et al., 2016). The presence of γ-H2A.X was investigated in protein extracts prepared from zeocin-treated plants and untreated controls, using immunodetection with specific γ-H2A.X antibody (Figure 6). As expected, a weak γ-H2A.X signal was observed in untreated protein extracts in all four lines (WT, *fas1*, *m123-2, fas1m123-2)* whereas zeocin treatment led to induction of γ-H2A.X and increased signal intensity (Figure 6). However, the intensity of γ-H2A.X signal, when compared to anti-H3 immunodetection used as a loading control, showed that phosphorylation of H2A.X in *fas1m123-2* was comparable to the other three lines, indicating that the first step in DNA damage signalling in *fas1m123-*2 progressed normally.

**Figure 6.**
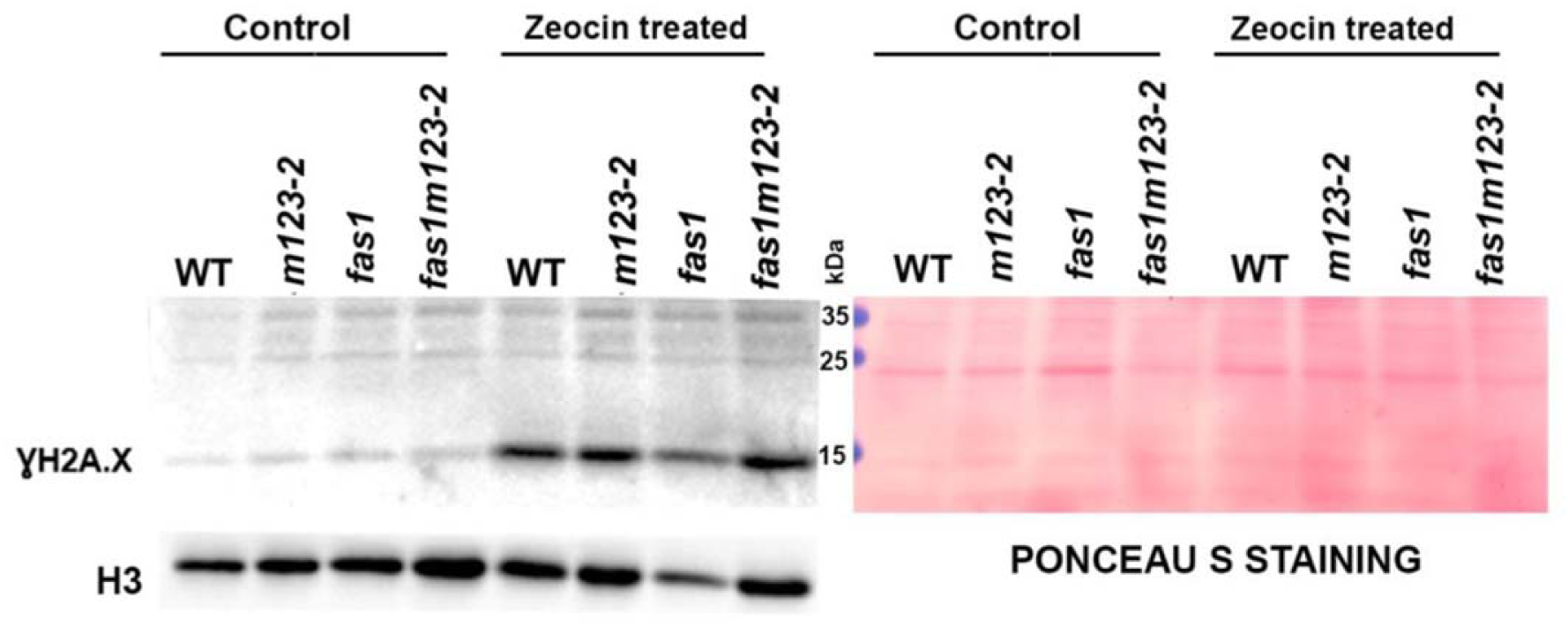
DNA damage signalling occurs normally in *fas1m123-2.* Western blot analysis of phosphorylated H2A.X (left panel) in untreated (left part) and zeocin-treated (right part) WT plants and *fas1, m123-1* and *fas1m123-2* mutants. Ponceau S staining (right panel) and histone H3 control blot are shown as loading controls.

### Stability of repetitive elements is enhanced by depletion of *NAP1* in *fas1* plants

A mutation in *FAS1* causes telomere shortening and the loss of 45S rDNA that occurs from the G1 generation onwards (Mozgova et al., 2010). Both repeats can be recovered when functional genes are introduced, but the recovery of 45S rDNA was shown to be much more stochastic than initially expected (Pavlistova et al., 2016). To elucidate the effect of simultaneous depletion of *NAP1* and *FAS1* on the amount of 45S rDNA, we used quantitative PCR. The 45S rDNA copy number was analysed in parental plants used for crossing (*fas1* G1 and *m123-2*, unknown generation), in heterozygous F1 and in four subsequent generations of *fas1m123-2* mutants (3 independent lines shown on Figure 7A and Supplemental Figures 3A and 4A). We also used *fas1* (G3) and another *m123-2* as controls. The parental *fas1* (G1) plants contained approx. 50% of the 45S rDNA, and *m123-2* had a similar number of copies of 45S rDNA as WT (Figure 7A). The arising F1 generation showed an occurrence of more than 60% of 45S rDNA copies when compared to WT. Interestingly, the absence of *NAP1* genes in *fas1* mutants suppressed the loss of 45S rDNA. Its level in *fas1m123-2* was maintained between 60-80% of the WT and this level became stable across generations. In the *fas1* (G3) control, in contrast, 45S rDNA reached as low as ∼30% of the copies in WT (Figure 7A), which further declined to 15-20% (Mozgova et al., 2010; Pontvianne et al., 2013).

**Figure 7.**
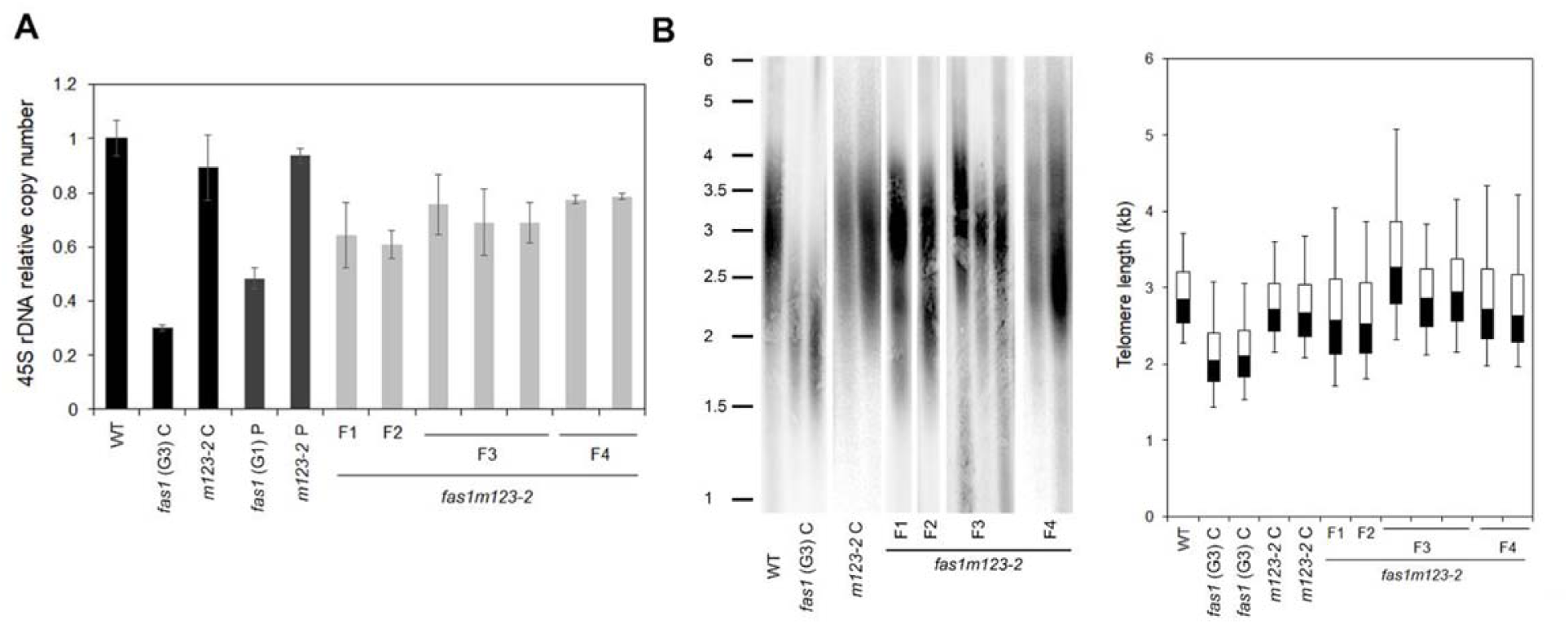
Telomere and 45S rDNA copy number in the *fas1m123-2* mutant return to WT levels. A) Q-PCR analysis of relative 45S rDNA copy number in WT, *fasl* (G3) C (control), *m123-2* C (control) and *fas1m123-2* mutant (F1 - F4 generations of line 5). Parental (P) plants *fas1* (G1) P and *m123-2* P are also shown. F1 is the heterozygous generation and F2 is the first homozygous generation of *fas1m123-2* mutant. Three biological replicates were used in *fas1* (G3) C, *m123-2C* and F3 and F4 of *faslm 123-2.* In the case of *fas1* P, *m 123-2* P and F1, F2 of *faslm 123-2* generations, 1 biological replica is shown. Error bars indicate standard deviations. B) Telomere lengths determined by terminal restriction fragment analysis in WT, *fas1* (G3) C (control), *m123-2C* (control) and *fas1m123-2* mutant (F1 - F4 generations of line 5). F1 is the heterozygous generation and F2 is the first homozygous generation of the *fas1m123-2* mutant. Graphical representation of telomere lengths was performed by evaluation of signal intensity from the membrane. Box charts show the first and third quartiles separated by the median value and SD intervals above and under the box representing the maximum and minimum of the telomere lengths.

NAP1;1-3 share the same molecular pathway with NRP1 and 2, particularly in the DNA damage response involving HR. We took advantage of the mutant line *fas2nrp1-*2 (Gao et al., 2012), where HR is significantly reduced compared to *fas2*, and we quantified the amount of 45S rDNA. In 3 subsequent generations of the *fas2nrp1-2* mutant and the control *fas2* mutant, we observed a progressive loss of 45S rDNA copies only in *fas2.* Although the levels of 45S rDNA were quite heterogeneous between *fas2nrp1-2* individuals, Supplemental Figure 5, the 45S rDNA repeats were protected against progressive loss in this line, similarly to *fas1m123-2*.

The impact of simultaneous depletion of *FAS1* and *NAP1;1-3* on maintenance of telomere length was explored in four subsequent generations of *fas1m123-2*. In contrast to *fas1* showing systematic telomere erosion (Mozgova et al., 2010), *fas1m123-2* revealed an overall trend towards telomere recovery (Figure 7B). Although telomeres showed inter-individual variability (1.8 - 2.5 kb) across generations, rather than a homogeneous telomere length profile (Figure 7B and Supplemental figures 3B and 4B), the progressive loss of telomeres, typical for *fas1,* was not observed in *fas1m123-2*.

### Chromatin in *fas1m123-2* is more compact

According to previous reports, chromatin in *fas1/2* contains lower levels of heterochromatin caused by an altered ratio of H3.1/H3.3 and lower levels of the repressive histone marks, e.g. H3K27me1 (Jacob et al., 2014; Duc et al., 2015; Otero et al., 2016; Wollmann et al., 2017). These chromatin changes contributed to the *fas1/2* phenotype. As concluded from the analysis in *fas1m123-2*, disruption of *NAP1* genes in *fas1* significantly improved overall plant growth and fitness, the response to genotoxic stress, and induced expression of DNA repair genes. To further test if these changes could be detected at the level of chromatin, we carried out digestion with Micrococcal nuclease (MNase) to determine chromatin accessibility in individual plant lines. The accessibility of chromatin to MNase digestion is determined by chromatin composition and structure, e.g. the proportion of different histone variants and other protein complexes (Fyodorov and Kadonaga, 2003; Chereji et al., 2017; Kubik et al., 2017). Nuclei prepared from leaves of WT, and *fas1, m123-2* and *fas1m123-2* mutants, were simultaneously digested with two different activities of MNase, optimised to achieve partial digestion.

After separation on an agarose gel and staining with ethidium bromide, we observed predominantly mononucleosome-sized fragments in the *fas1* sample, while in the *fas1m123-2*, mono-, di-, tri-nucleosomes were detected, similarly to the WT and *m123-2* (Figure 8A and Supplemental Figure 6A). Occurrence of nucleosome-sized fragments was further analysed using the fragment analyser (Figure 8D). The peak of mononucleosomes was lower in *fas1* than in *fas1m123-2,* WT or *m123-2,* indicating that some DNA fraction in *fas1* was degraded by exonucleolytic activity. In *fas1m123-2,* higher levels of mono- and di-nucleosomal DNA fragments were detected. The concentration of DNA, as determined by Qubit, was comparable between individual samples (Supplemental Table 1). To extract information about chromatin organization of repetitive sequences affected in *fas1/2* mutants, we hybridised the gel-separated fragments with a selected set of probes. Two membranes were prepared, one containing an equal amount of DNA in each line, and the second with an enriched amount of DNA in *fas1* (to facilitate the detection of a reduced level of 45S rDNA in *fas1*). Hybridisation with bulk genomic DNA, 45S rDNA as well as with a telomeric probe revealed profound differences between *fas1* and *fas1m123-*2 (Figure 8B, C and Supplemental figure 6 B, C) Chromatin in *fas1* was extensively digested and 45S rDNA and telomeres accumulated in the mononucleosomal fraction. We used the 6^th^ generation of *fas1* plants, where only active and presumably a greater MNase sensitive fraction of 45S rDNA occurs and telomeres are shortened to ∼ 1.5kb. In contrast, in *fas1m123-2*, sensitivity to MNase decreased and the level of chromatin digestion seemed even lower than in WT or *m123-2*.

**Figure 8.**
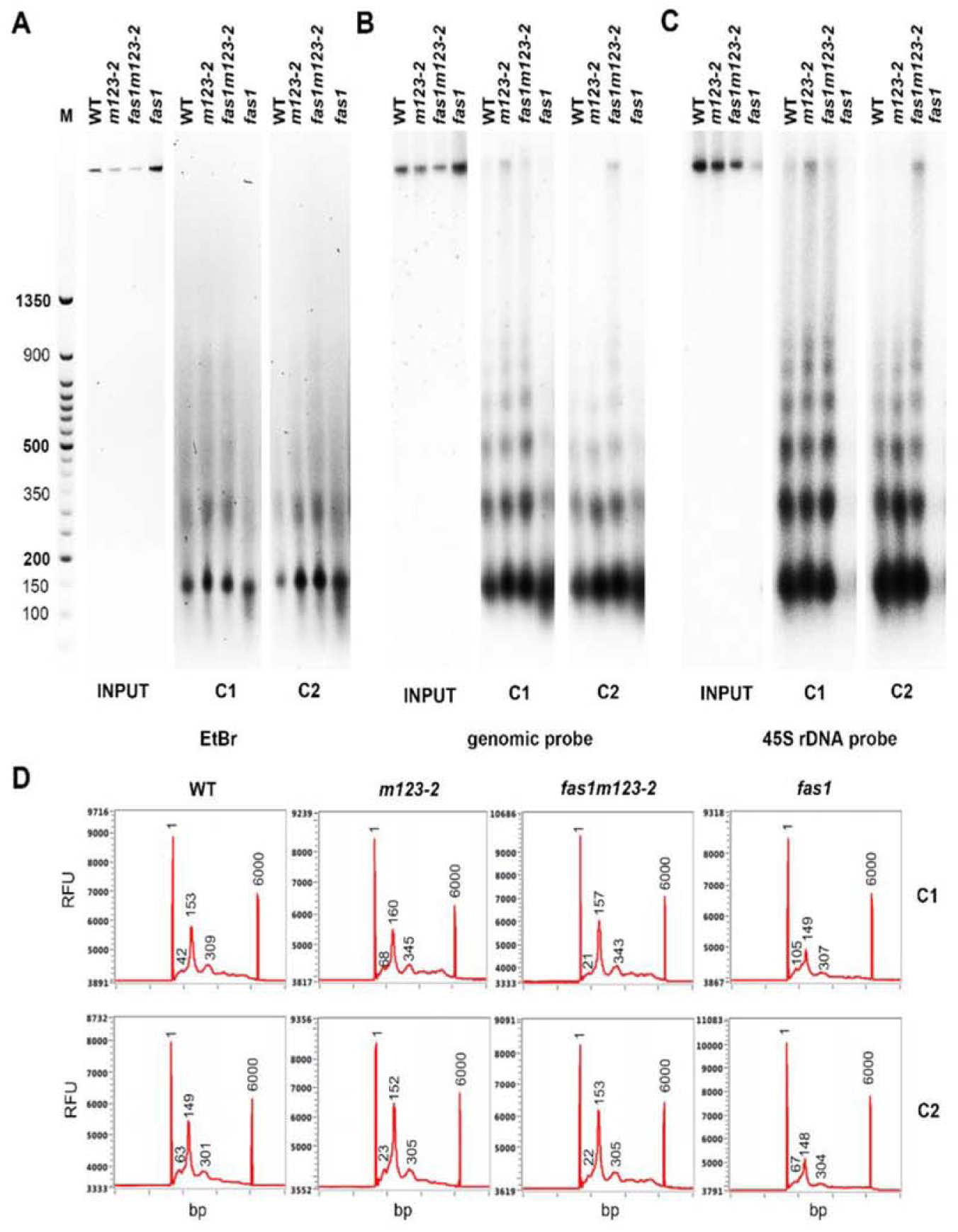
Chromatin in *fas1m123-2 is* protected against MNase. Chromatin prepared using 8-w-old leaves was digested with two different concentrations ol MNase-C1 corresponds to 80 Gel units and C2 corresponds to 160 Gel units. DNA fragments were separated on agarose gel or on the fragment analyser. M represents the molecular weight marker, INPUT and digested samples WT, *fas1, m123-2* and *fas1m123-2* are shown. A) Agarose gel stained with ethidium bromide 0.5 μg/ml. B) Hybridisation pattern using radioactiveiy labelled probe of bulk genomic DNA, Non-digested INPUTS represent 1/10 of the total DNA used for MNase digestion. C) Hybridisation pattern using radioactiveiy labelled probe specific for 45S ribosomal gene locus. Non-digested INPUTS represent 1/10 of the total DNA used for MNase digestion. D) Digested DNA separated on the Fragment analyser. Fragments of mono- and di-nucleosomes are shown. RFU represent relative fluorescence units.

This data confirmed our hypothesis that the improvement of phenotype occurring in *fas1m123-2* emerged as a consequence of global changes in chromatin structure.

### FAS1 and NAP1;1-3 proteins do not interact directly

Some histone chaperones that participate in the same molecular pathway mediate their functions via physical contact. CAF-1 complex, for example, directly interacts with an H3.3/H3.1 histone chaperone ASF1 (Tyler et al., 2001; Mello et al., 2002). We show here that remarkable genetic interaction occurs between H3/H4 and H2A/H2B histone cnaperone genes *fasi* and *napi;i-3,* respectively and we Turther asked whether FAS1 and napi;i-3 could form functional protein complexes *in vivo. We* thus employed bimolecular fluorescence complementation (BiFC), Figure 9A, and the yeast two hybrid assay (Y2H), Figure 9B, to test it. These experiments, however, did not prove any direct physical interaction between NAP1;1-3 and FAS1 chaperones, suggesting that cooperation between FAS1 and NAP1;1-3 occurs in another way. To exclude that the absence of interaction is caused by insufficient expression of FAS1 and NAP1;1-3 proteins, we employed two positive controls: conserved interactors FAS1 and ASF1 (Tyler et al., 2001; Mello et al., 2002) and NAP1;1/NAP1;1, known to form homodimers in plants (Liu et al., 2009). The positive controls worked as expected in Y2H (Supplemental Figure 7) confirming the reliability of our observations. BiFC confirmed the FAS1-ASF1 interaction *in vivo* and was localised in the nucleus (Supplemental Figure 7). NAP1;1 protein was localised in the cytoplasm in plants and its homodimers were formed in the same compartment (Supplemental Figure 7). For negative controls, we used the empty vectors/NAP1;1 and NAP1;1/NRP1 that did not interact in Y2H (Liu et al., 2009).

**Figure 9.**
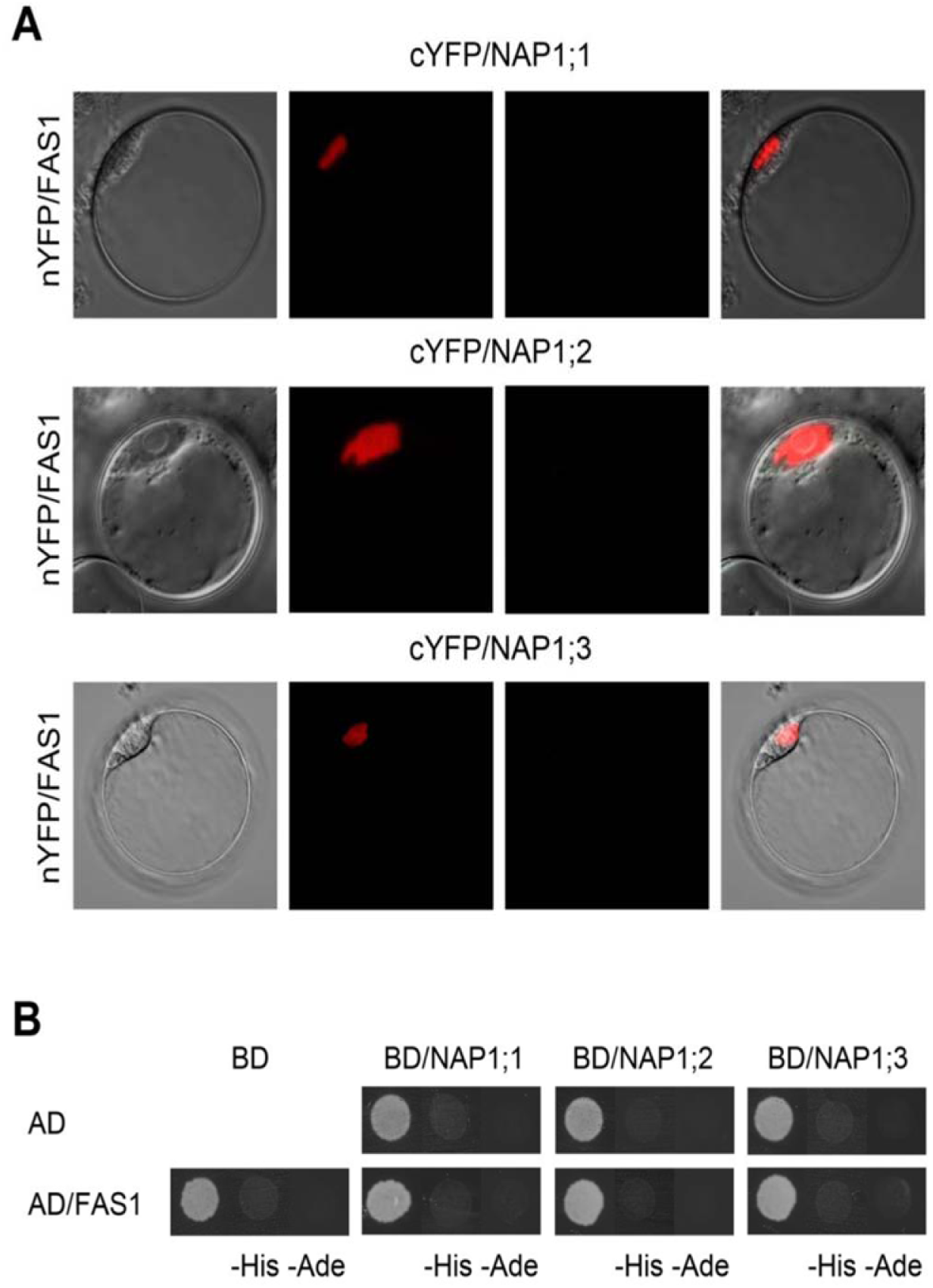
FAS1 does not interact with members of the NAP1 family proteins. A) Yeast two hybrid system. NAP1;1-3 and FASH in either pGADT7 or pGBKT7 expression vectors were co-transformed into the yeast strain PJ69-4a. Yeast cells were plated onto control medium (without Leu and Trp) and onto selective media lacking Leu, Trp and His (-His) and Leu, Trp, His and Ade (-Ade, strong interaction). The growth of yeasts on selective media indicates positive protein-protein interactions. B) Bimolecular Fluorescence Complementation. *Arabidopsis* protoplasts were co-transfected with nYFP/FAS1 and cYFP/NAP1;1 or cYFP/NAP1:2 or cYFP/NAP1;3 plasmids. Bright field, nuclear localization signal mRFP-NLS (red), the YFP fluorescence (green) and the combined images were visualized under a confocal microscope 16 h after transfection. The scale bar represents 10 μm. Experiments were repeated three times with similar results.

## DISCUSSION

Histone chaperones are involved in numerous protein complexes regulating the balanced composition of nucleosomes, a precise combination of activating or repressive histone marks, and further determining individual chromatin states, e.g. (Roudier et al., 2011; Sequeira-Mendes et al., 2014; Hammond et al., 2017).

We have focused here on the functional interaction between the major plant H3.1 chaperone CAF-1, and members of the NAP1-protein group that are known to bind H2A/H2B histones (Liu, 2009, Zhou 2016). CAF-1 deficient plants are viable, but the mutation causes progressive growth defects, finally leading to retarded infertile plants (Mozgova et al., 2010; Mozgova et al., 2018). The typical *fas1* mutant phenotype is pleiotropic and the most obvious defects include fasciated stems and dentate leaves (Exner et al., 2006; Mozgova et al., 2010; Varas et al., 2015). The disruption of *NAP1* genes, on the other hand, has no obvious phenotype at the macroscopic level (Liu et al., 2009). At the subcellular level it causes decreased HR and increased sensitivity to UVC radiation (Liu et al., 2009; Gao et al., 2012).

We based this study on the intriguing observation that the quadruple mutant *fas1m123-2* appears phenotypically as WT rather than *fas1* (Figure 1). A further investigation of this mutant line revealed that most of the subcellular phenotypes found in *fas1* were abrogated in *fas1m123-2*. This unexpected genetic interaction between two histone chaperones suggests that many of the abnormalities occurring in *fas1* plants are mediated via *NAP1* molecular pathways when CAF-1 is dysfunctional. Although *fas1* is a well described mutation, the primary cause of the *fas1* phenotype has not been fully explained and its dependence on NAP1 proteins was not postulated. This study unmasks a so far unrecognised function of *NAP1* in the maintenance of chromatin structure, suggesting that NAP1 proteins are required for the maintenance of the open chromatin state in *fas1,* as summarised in Figure 10.

**Figure 10.**
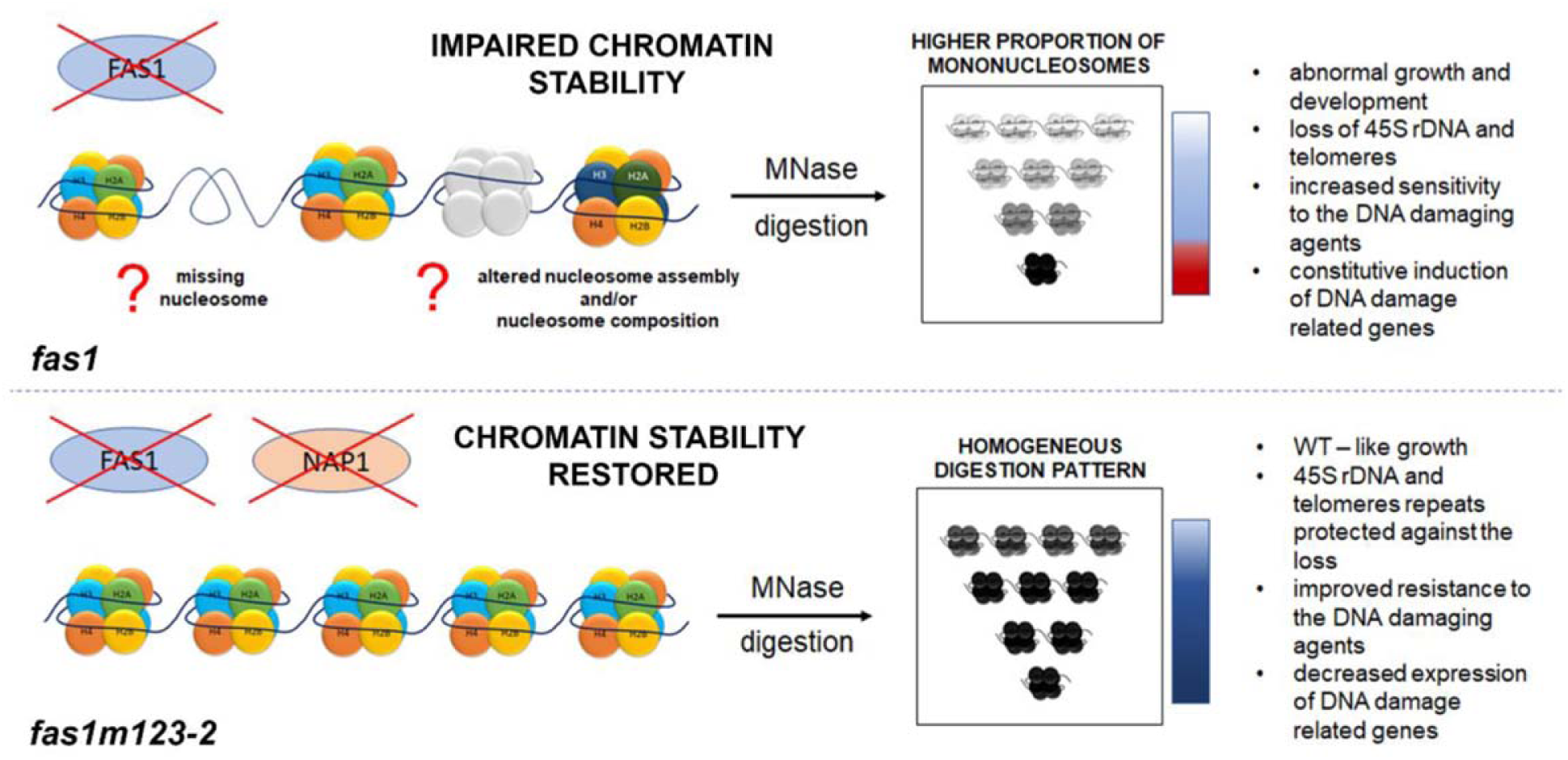
Overview of the contribution of *NAP1* to the *fas1* phenotype. Increased chromatin accessibility observer in *fas1* becomes restored after disruption of *NAP1* genes. additionally, many ot the *fas1* defects are improved in *fas1m123-2*.

### Maintenance of chromatin stability in *fas1m123-2*

CAF-1 is an H3.1 histone chaperone and its deficiency can, at least partially, be compensated for by *HIRA* or *ATRX* complementary pathways, as shown recently (Duc et al., 2015; Duc et al., 2017). This compensation, however, affects the ratio of H3.1/H3.3, otherwise important for correct chromatin assembly and the epigenetic set up during early development (Otero et al., 2016; Benoit et al., 2019). The altered H3.1 loading and H3.1/H3.3 imbalance, as observed in *fas1,* relates to the: i) lower abundance of nucleosomes, particularly in non-transcribed regions (Munoz-Viana et al., 2017); ii) decreased levels of H3K27me1 histone mark (Jacob et al., 2014; Wollmann et al., 2017); iii) instability of 45S rDNA and telomere repeats and decondensation of chromocenters (Kirik et al., 2006; Mozgova et al., 2010; Varas et al., 2015); and/or iv) changes in the transcription level of a selected set of genes (Bouveret et al., 2006; Ramirez-Parra and Gutierrez, 2007; Mozgova et al., 2015). We looked at some of these deleterious effects of the *fas1* mutation in *fas1m123-*2 and found that many changes were suppressed when *NAP1;1-3* genes were disrupted. We showed that chromatin in *fas1m123-2* became more protected against MNase treatment in bulk DNA, telomeres, as well as in 45S rDNA compared to *fas1* plants, and that protection was even higher than in WT (Figure 8B, C, Supplemental Figure 6C). MNase is known to more rapidly digest the A/T rich and nucleosome-free regions, promoters occupied by non-nucleosomal complexes, or nucleosomes enriched in certain histone variants, e.g. H3.3 or H2A.Z (Jin et al., 2009; Chereji et al., 2017). The increased digestion rate in *fas1* may reflect the lower density of nucleosomes and the different H3.1/H3.3 balance, as previously observed in *fas1* (Kirik et al., 2006; Otero et al., 2016; Varas et al., 2017), thus the higher protection of chromatin in *fas1m132-2* possibly suggests changes in chromatin structure in the quadruple mutant (Figure 10). The total abundance of H3 in *fas1* or *fas1m123-2* did not differ (Figure 8), indicating that the observed changes were not caused by altered levels of H3. Moreover, NAP1 is considered to be an H2A/H2B histone chaperone in plants and such an alteration could be expected at the level of H2 turnover or even other protein complexes. An interesting candidate could be the H2A.W variant, important for heterochromatin formation and internal DNA damage repair (Yelagandula et al., 2014), whose levels were shown to drop in *fas1* (Benoit et al., 2019). Moreover, we show that individual *NAP1* genes are upregulated in *fas1,* potentially causing an imbalance during chromatin assembly in *fas1*.

### Repetitive elements demonstrate improved genome stability in *fas1m123-2*

Destabilisation of repetitive elements often represents a threat to genome integrity (Sfeir et al., 2009; Ivics and Izsvak, 2010; Levine et al., 2016; Ozer and Hickson, 2018). Mutations in chromatin components and DNA repair genes often affect the stability of 45S rDNA genes and telomeres in *Arabidopsis* (Vannier et al., 2009; Schrumpfova et al., 2011; Rohrig et al., 2016; Dorn et al., 2019) and disruption of *FAS1/2*, in particular, shows one of the strongest effects (Mozgova et al., 2010; Pontvianne et al., 2013). We found that in *fas1m123-2* plants, the loss of 45S rDNA as well as telomeres was largely supressed when compared to *fas1* (Figure 7, Supplemental Figures 3 and 4).

45S rDNA clusters are considered as fragile sites and potential targets for recombination (Dvorackova et al., 2015; Huang et al., 2018). Homologous recombination occurs naturally, particularly inside 45S rDNA clusters and contributes to the maintenance of copy number (Kobayashi et al., 2004). *NAP1;1-3* together with *NRP1* and *NRP2* participate in HR (Gao et al., 2012; Zhou et al., 2016) and their deficiency in *nrp1-2* mutants causes HR downregulation. *fas1/2* plants, in contrast, show elevated levels of recombination activity (Kirik et al., 2006) and this declines significantly when *FAS2* and *NRP1-2* are disrupted in *fas2nrp1-2* (Gao et al., 2012).

In *fas2nrp1-2* plants, recombination activity remains at WT levels and, as we observed, 45S rDNA was partially protected against the loss (Supplemental Figure 5). A similar situation occurs in *fas1/2rad51B,* where somatic HR factor RAD51B is missing (Muchova et al., 2015). Considering the functions of NAP1 and NRP1-2 in HR, we propose that the efficiency of HR might be lower in *fas1m123-2* plants, similarly to *fas2nrp1-2,* and that it contributes to the stabilisation of 45S rDNA repeats.

While homologous recombination occurs naturally within 45S rDNA clusters, it is suppressed at telomeres (Vannier et al., 2009; Eckert-Boulet and Lisby, 2010). Consistently, telomeres are not affected by changes in the efficiency of HR, and short telomere phenotypes still occur in *fas2nrp1-2* as well as in *fas1/2rad51b* (Gao et al., 2012; Muchova et al., 2015). The observed suppression of telomere loss in *fas1m123-2* suggests the contribution of NAP1;1-3 in the telomere erosion process in *fas1*. The underlying mechanism however, remains to be elucidated.

### NAP1 proteins participate in root development

We show here that the short-root phenotype typical for *fas1* did not occur in *fas1m123-2* (Figure 3). Root growth and development are regulated by a set of genes whose transcript levels are deregulated in *fas1/2* (Bouveret et al., 2006; Exner et al., 2006). FAS2 and NRP1-2 act cooperatively in SCN maintenance and the short-root phenotype was strongly aggravated in the combined mutant *fas2nrp1-2* (Kaya et al., 2001; Zhu et al., 2006; Huang et al., 2018). The roots of *fas1m123-2* grow as well as WT. One plausible explanation is that NAP1;1-3 act upstream of FAS1-2 and NRP1-2, and that their disruption in *fas1m123-*2 affects the transcription of root-specific genes leading to improved root growth and patterning. Alternatively, different genetic interactions could occur when either FAS1 or FAS2 are considered. Future studies are still necessary to investigate more precisely the detailed molecular interactions between FAS1-2, NRP1-2 and NAP1;1-3 in the regulation of SCN.

### Altered expression of DNA repair genes in *fas1m123-2* reflects DNA damage signalling and cell cycle progression

The lower occurrence of SCN cell death in *fas1m123-2* points to the importance of *NAP1;1-3* in the DNA damage response pathway in *fas1* and explains why *fas1m132-2* plants acquired tolerance to genotoxic agents. The lack of root cell death was previously shown in *fas1atm* mutants, for example, the line deficient in the important DNA damage kinase ATM (Hisanaga et al., 2013). Our view is further supported by changes in two DNA damage marker genes *BRCA1* and *PARP1* in a quadruple mutant. Although the expression of *BRCA1* and *PARP1* in *fas1m123-2* was 3-fold higher than in the WT, it was lower than in *fas1* (Figure 5). Interestingly, the observed loss of *BRCA1* and *PARP1* induction after CPT treatment in *fas1* also occurred in *fas1m123-2* plants. It is in line with the previous observations in *fas1* (Bouveret et al., 2006; Hisanaga et al., 2013; Huang et al., 2018) and it indicates that the pathway responsible for induction of both genes was not fully rescued by disruption of *NAP1;1-3* genes in *fas1m123-2*.

Enhanced expression of *BRCA1* and *PARP1* in *fas1* is dependent on ATM, leading to the activation of the G2/M checkpoint (Hisanaga et al., 2013). Altered cell cycle progression often relates to retarded plant growth that in the case of *fas1*, is compensated by fewer large cells with higher levels of ploidy (Endo et al., 2006; Ramirez-Parra and Gutierrez, 2007). When ATM kinase is deleted in the *fas1atm* mutant, expression of *BRCA1* and *PARP1* returns to WT levels and developmental defects, including a reduction in cell number and a high ploidy phenotype, are partially rescued (Hisanaga et al., 2013). The fertility of *fas1atm* plants, however, remains poor.

We show here that the effect of *NAP1;1-3* disruption on the *fas1* phenotype resembles the effects of ATM deficiency: the ploidy phenotype was rescued in *fas1m123-2* (Figure 1D), moreover; *fas1m123-2* plants were fully fertile (Figure1B, C).

Altogether we propose that a more efficient DNA repair process is activated in *fas1* in the absence of NAP1 functions. *NAP1* genes act either upstream of *FAS1*, or they act as key mediators of DNA damage response in *fas1*. Which step in the DNA damage pathway is affected is not yet clear. We showed that H2A.X phosphorylation, indicating activation of *ATM,* was not affected in *fas1m123-2.* The DNA damage sites are thus still present in *fas1m123-2,* and *NAP1* is not required to activate *ATM* in *fas1*.

This work presents another step in an investigation of interrelated processes of chromatin assembly and maintenance of genome stability/DNA damage repair. Our observations contribute to the understanding of the function of NAP1;1-3 histone chaperones in plants and describes genetic interactions between *NAP1;1-3* and *FAS1* that have not yet been shown in any other model. Understanding of how *fas1m123-2* plants cope with a deficiency in two important histone chaperones represents a complex scientific question, potentially important even outside of the plant kingdom. We conclude that in *fas1m123-2* some alternative histone chaperone pathway is activated, efficiently overcoming both - CAF-1 and NAP1 dysfunctions.

## METHODS

### Plant material and growth conditions

In this study we used previously described mutant lines of *A. thaliana,* which were derived from the Columbia ecotype: *fas1-4* (NASC: N828822, SAIL_662_D10 (Exner et al., 2006; Mozgova et al., 2010), *NAP1* loss-of-function mutant *m123-2* (*nap1;1, nap1;2, nap1;3* alleles correspond respectively to SALK_013610, SAIL_373_H11 and SALK_131746 of T-DNA insertion lines from ABRC) (Liu et al., 2009; Zhou et al., 2016). Quadruple mutants were obtained in our laboratory by crossing the first generation of *fas1-4* plants with undefined generations of *m123-2* plants provided by Wen-Hui Shen (CNRS, Strasbourg, France and Fudan University, Shanghai, China). Mutant lines were segregated from heterozygous plants obtained in the F1 generation). Three independent lines (5, 20 and 22) were obtained and then propagated into the F3 consecutive generation.

All seeds were first surface sterilized (70% ethanol/10 min, 99% ethanol/5 min) and plated on half-strength agar Murashige and Skoog medium (½ MS medium) with 1% sucrose. After 2 days of stratification (4°C/dark) plates were transferred to the growth chamber and pre-grown for 2 weeks under long day (LD) conditions (16 h light - 21°C/8 h dark −19°C/ 50-60% relative humidity). Seedlings were collected directly from plates or transferred into soil after 2-weeks germination and were grown under the same long day conditions.

### DNA isolation and plant genotyping

DNA was isolated from 5-w-old rosette leaves using the Dellaporta protocol (Dellaporta et al., 1983). Concentration and integrity of DNA was determined using a 1% (w/v) agarose gel containing 0.5 μg/mL EtBr and a Gene Ruler 1 kb DNA ladder (Fermentas) as a standard and Multi Gauge software (Fujifilm) for evaluation.

For genotyping, DNA was isolated using GenElute Plant Genomic DNA Miniprep Kits (Sigma), and 1 μL of DNA was used in a PCR reaction using 0.3 μΜ primers and 5U of MyTaq DNA polymerase (Bioline).

### Flow Cytometry

Leaves of 22-d-old plants, grown for 10 days under LD and for 12 days under SD conditions (10 h light - 21°C/14 h dark - 19°C/ 50-60% relative humidity), were chopped in Galbraith buffer containing 0.3 % of Triton (Galbraith et al., 2011), filtered through the 30 μm Partec filter and stained with 100 μg/mL of PI and 100 μg/mL of RNase A.

### Quantitative PCR (Q-PCR)

For relative quantification of 45S rDNA, gDNA from 5-w-old rosette leaves was used. Where possible, three biological replicas were used (DNA isolated from leaves of 3 different plants). Q-PCR was performed according to standard procedures using FastStart SYBR Green Master (Roche) and a combination of 18SFw and 18SR primers for 18S rDNA (226 bp long product) normalized to UBIQUITIN 10 (see Supplemental Table 2). The analysis was performed by StepOnePlus Real Time PCR system (Applied Biosystems) under the following conditions: 95°C/7 min; 40 cycles of 95°C/30 sec, 56°C/30 sec, 72°C/30 sec and 72°C/5 min followed by standard melting analysis. Reactions for each sample were carried out in technical triplicates in three independent Q-PCR experiments and the mean value was counted. Individual plants are shown, thus standard deviation relates to variation in the technical replicates.

For gene expression studies, total RNA was extracted from seedlings using NucleoSpin® RNA Plant kit (MACHEREY-NAGEL) according to the manufacturer’s instructions. Isolated RNA was treated with Turbo DNA-free (Ambion) for 2 x 30 min to eliminate contaminants of the DNA. RNA concentration was determined using a NanoDrop 2000c (Thermo Scientific) and RNA integrity was checked on a 1% (w/v) agarose gel containing 0.5 μg/mL EtBr.

Synthesis of cDNA was performed using 1 μg of RNA, random nonamers (Sigma) or oligo-dT primers (Thermo Fisher Scientific) and an M-MuLV reverse transcriptase kit (Finnzymes). Q-PCR was performed in triplicates for all the samples to analyse expression of *NAP1*;*1, NAP1*;*2*, *NAP1*;*3*, *PARP1* and *BRCA1* genes (primers listed in Supplemental Table 2) with normalization to UBIQUITIN 10 expression under the following conditions: 95°C/7 min; 30 cycles of 95°C/30 sec, 54 - 56°C (depending on the gene)/30 sec, 72°C/30 sec and 72°C/5 min followed by standard melting analysis. Used biological replicates represent 3 independent pools of seedlings or leaves from 3 independent plants, SD reflects the variability between biological replicas.

### Terminal Restriction Fragment Analysis (TRF)

For TRF analysis, gDNA extracted from 5-w-old rosette leaves was used. Seven hundred nanograms of gDNA were digested overnight using 10 U of *Mse*I (NEB). TRF products were separated on a 0.8% agarose gel containing 0.5 μg/mL EtBr at 1.5 V/cm for 16 hours at room temperature. Agarose gels with TRFs were blotted in 0.4 M NaOH onto Hybond XL membrane (GE Healthcare). TRFs were hybridized with radioactively [Y-^32^P]-dATP labelled TEL C 5‘-(CCCTAAA)_4_-3 overnight at 55°C. Alternatively, plant telomeric concatemers labelled with [a-^32^P]-dATP were used and hybridisation was performed overnight at 65°C. For more details see also (Neplechova et al., 2005). TRF signals were visualized using the FLA 9500 (Fuji Film). For the evaluation of visualised telomere fragments we used WALTER toolset, an upgraded version of previous evaluation tool developed in our laboratory (Zachova et al., 2013). As a molecular weight standard, we used the Gene Ruler 1 kb DNA ladder (Fermentas). The length of telomeres is presented using a box-and-whisker plot where the bottom and the top of the box represent the first and third quartiles, separated by the median. Standard deviation (SD) intervals above and under the box represent the minimum and maximum of telomere lengths.

### Sensitivity test

Genotoxic effects of selected genotoxic agents were evaluated on plants either grown on the agar plates or incubated overnight in liquid MS medium, both containing an appropriate genotoxic agent. Seeds were surface sterilized with ethanol and transferred onto the ½ MS agar with 1% sucrose, stratified (2 days at 4°C in dark) and grown under long day (16h/8h) conditions. After 4 days, seedlings were transferred onto control plates (½ MS agar medium + 1% sucrose) or plates supplemented with genotoxic agents (10 nM CPT, 1 μg/mL bleomycin, 20 μg/mL zeocin, 1 μΜ MMC or 0.125 mM MMS). Seedlings were grown for the next 6 days under long day conditions, documented and collected for RNA isolation. Root lengths of at least 20 plants in three experiments were measured using the ImageJ software. Alternatively, PI staining was applied to determine the level of apoptosis in the root stem cell niche. Ten-d-old CPT treated seedlings as well as 4-d-old seedlings treated with CPT overnight were used in the PI assay.

Propidium iodide staining was performed by incubation of root tips for 1 min in 10 μg/mL PI/water solution (Fulcher and Sablowski, 2009), followed by visualisation of the root tips by laser scanning confocal microscopy (Leica SPE).

### Alexander staining

For the detection of pollen grain viability, a modified Alexander staining protocol lacking toxic agents was used (Peterson et al., 2010). Flower buds from ∼3 month old plants were collected and fixed in ethanol:glacial acetic acid 3:1. After 2 hours, the solution was applied directly to the microscope slide containing excised anthers. The number and colour of pollen grains (red - viable; green - non-viable) were evaluated using a Zeiss Axioscope microscope with digital camera.

### Cloning

Entry clones containing complete CDS sequences of *NAP1;1*, *NAP1*;2, *NAP1*;3, *FAS1*, *FAS2*, *NRP1* and *ASF1a* genes were obtained by cloning using PCR and the Gateway technology (Invitrogen). cDNA was synthesized from total RNA of 10-d-old seedlings using M-MuLV reverse transcriptase (New England Biolabs); each gene was amplified using specific primers (see Supplemental Table 2) and Phusion HF DNA polymerase (Finnzymes). For cloning, Gateway compatible vectors pDONR207, pGADT7 and pGBKT7 for Y2H and pSAT1-nEYFP and pSAT1-cEYFP for BiFC were used and cloned using BP and LR clonases, (Invitrogen). Plasmid DNA was isolated with QIAprep Spin Miniprep Kit (QIAGEN) for the Y2H assay or Nuclebond® Xtra Midi (MACHERE-NAGEL) for BiFC. All procedures were carried out in accordance with manufacturer’s protocols.

### Yeast two-hybrid (Y2H) system

PJ69-4α strain of *Saccharomyces cerevisiae* was used for the expression of proteins of interest using the Matchmaker Gal4-based yeast two hybrid system (Clontech). Tested bait/prey combinations were co-transformed into PJ69-4α and cultivated at 28°C. Positive interactions were tested on plates with SD agar lacking Leu, Trp and His and strong interactions on plates lacking Leu, Trp, His and Ade. Histidine-lacking plates contained increasing concentrations of 3-aminotriazol (3-AT), which inhibits His3 activity and correlates with the higher binding affinity of the proteins. For the control of auto­activation, co-transformations with empty vectors were tested. Each test was carried out in three repeat experiments.

### Bimolecular fluorescence complementation (BiFC)

For the BiFC experiment, protoplasts from root tips of 7-d-old seedlings grown on ½ MS medium under long day conditions were used. For releasing the protoplasts from root tips, an enzyme solution (0.4% Cellulase, 0.2% Macerozyme, 0.4 M mannitol, 20 mM KCl, 20 mM MES with pH6, 10 mM CaCl_2_, 0.1% BSA) was used and 1-2 mm long pieces of roots were incubated for 2-5 h/100 rpm at RT. Protoplasts were then filtered through a 70 μm cell strainer and 2x washed with W5 solution (154 mM NaCl, 125 mM CaCl_2_, 5 mM KCl, 2 mM MES (pH 6), 5 mM glucose) (4°C, 200 g, 4 min). Protoplasts were counted and resuspended in a certain volume of MMg solution (0.4 M mannitol, 15 mM MgCl_2_, 4 mM MES (pH 6)) to achieve a protoplast concentration of 300,000 cells per mL. 10 μg of each plasmid DNA in a final volume of 20 μL were used and 200 μL of protoplasts in MMg solution and 220 μL of PEG solution (40% PEG, 0.2 M mannitol, 0.1 M CaCl_2_) were added to the mixture and incubated (20 min, 50 rpm, RT). To quantify transformation efficiency, the protoplasts were co-transfected with the plasmid expressing mRFP and the nuclear localization signal of the VirD2 protein of *A. tumefaciens* (mRFP-VirD2NLS) (Citovsky et al., 2006). After 20 min incubation, protoplasts were washed in W5 solution (4°C, 4 min, 200 g), resuspended in 600 μL of W5, added to a tissue culture plate and incubated overnight in the dark at RT. The next day, fluorescence was observed using a Zeiss LSM 8000 laser scanning confocal microscope with YFP (Alexa Fluor 488) and RFP (Texas Red) filters.

### Mnase assay

Chromatin isolation was performed according to (Munoz-Viana et al., 2017), using 400 mg of leaves of 5-8-w-old plants. Two different concentrations (80 and 160 Gel Units) of MNase (NEB) were used. To quantify nucleosome size, precipitated DNA was analyzed with a fragment analyzer (Agilent, CA, USA) using the high sensitivity Next Generation S equencing kit 1-6000 (DNF-474) and the concentrations were measured on a Qubit Fluorometric Quantification system (Thermo Fisher Scientific), both according to manufaturer’s instructions. Nuclesome size was evaluated with the AATI ProSize software. DNA fragments were then separated on a 2% agarose gel containing 0.5 μL/mL of ethidium bromide, transferred onto a Hybond XL membrane (GE Healthcare) and subjected to hybridization with radioactively labeled sequence-specific probes: cloned fragments of 18S+25S rDNA (Mozgova et al., 2010) or plant telomeric concatemers, see above. The template for the bulk DNA probe was generated using isolated nuclei (Galbraith et al., 2011) and GenElute™ Plant Genomic DNA Miniprep Kit (Sigma).

### Western blot and immunodetection of proteins

Seven-d-old seedlings (∼ 0.5 g) were incubated either in liquid ½ MS or in ½ MS containing 100 μg/mL of zeocin (Invitrogen) for 2.5 h. Seedlings were then quickly dried on the filter paper and nuclei were released using an homogeniser (Quiagen) and Galbraith buffer (Galbraith et al., 2011) supplemented with protease and phosphatase inhibitors (Serva). Isolated nuclei were spun down for 10 min at 1500g, 4°C and resuspended in approx. 300 μL of Galbraith buffer containing both inhibitors. Protein concentration was determined by Bradford assay. Lysis was performed in SDS loading buffer (250 mM Tris-Cl, pH 6.8, 4% (w/v) SDS, 0.2% (w/v) bromophenol blue, 20% (v/v) glycerol, 200 mM β-mercaptoethanol) at 80°C Tor 10 min. Proteins were analysed by 12.5% SDS-PAGE and transferred onto the membrane at 36 mA overnight and 4°C (Schrumpfova et al., 2014). Equal gel loading was checked by Ponceau S staining (0.1% (w/v) Ponceau S in 5% (v/v) acetic acid). Membranes were blocked with 5% non-fat dry milk in Tris-buffered saline/Tween, and probed using anti-H3 (ab1791, Abcam) or anti YH2A.X (Sigma H5912), 1:1000. Signals were detected using secondary polyclonal horseradish peroxidase-conjugated antibody (Sigma A0545), 1:2000. Immunoreactive bands were visualized using LumiGLO reagent and peroxide (Cell Signaling Technology, http://www.cellsignal.com) on a Fusion CCD system.

## ACCESSION NUMBERS

**See Supplemental Table 3.**

## SUPPLEMENTAL DATA

**Supplemental Figure 1.** Ploidy levels in WT, *fas1*, *m123-2* and *fas1m123-2*.

**Supplemental Figure 2.** Sensitivity of WT, *fas1*, *m123-2* and *fas1m123-2* plants to DNA damaging agents.

**Supplemental Figure 3.** Relative 45S rDNA copy number and telomere length in line 20 of the *fas1m123-2* mutant.

**Supplemental Figure 4.** Relative 45S rDNA copy number and telomere length in line 22 of the *fas1m123-2* mutant.

**Supplemental Figure 5.** 45S rDNA is protected against loss in the *fas2nrp1-2* mutant.

**Supplemental Figure 6.** Chromatin in *fas1m123-2* is protected against MNase digestion.

**Supplemental Figure 7.** Protein-protein interactions revealed by BiFC and Y2H assays.

**Supplemental Table 1.** Measurement of DNA extracted in INPUTs and Mnase digested samples.

**Supplemental Table 2.** List of primers.

**Supplemental Table 3.** Accession numbers.

## ACKNOWLEDGEMENTS

Core Facilities: CELLIM of CEITEC supported by MEYS 279 CR (LM2015062 Czech-BioImaging), Genomics supported by NCMG research and Plant Sciences of CEITEC MU are gratefully acknowledged for obtaining the scientific data presented in this paper.

We would like to thank J. Kapustova and M. Chropovska for excellent technical support and prof. Shen (CNRS, Strasbourg, France and Fudan University, Shanghai, China) for providing us with necessary material.

Funding: This work was supported by the Czech Science Foundation (projects 19-11880Y and 17-09644S); by Ministry of Education, Youth and Sports of the Czech Republic projects CEITEC 2020 (LQ1601) and INTER-COST (LTC18048) and by COST INDEPTH CA16212.

## AUTHOR CONTRIBUTIONS

MD and ES provided ideas for the project and obtained funding. MD, MND and KK contributed to the concept of this work; KK and MND performed the majority of the experimental work and data analysis. KK prepared the figures. MD and MND wrote the manuscript with input from KK. ES critically revised the MS. TL performed the flow cytometry.

## Parsed Citations

Almouzni, G. (2009). Chromatin assembly factors and the challenges of DNA replication and repair. Faseb J 23.

Andrews, A.J., Downing, G., Brown, K., Park, Y.J., and Luger, K. (2008). A thermodynamic model for Nap1-histone interactions. The Journal of biological chemistry 283, 32412–32418.

Andrews, A.J., Chen, X., Zevin, A., Stargell, L.A., and Luger, K. (2010). The histone chaperone Nap1 promotes nucleosome assembly by eliminating nonnucleosomal histone DNA interactions. Molecular cell 37, 834–842.

Benoit, M., Simon, L., Desset, S., Duc, C., Cotterell, S., Poulet, A., Le Goff, S., Tatout, C., and Probst, A.V. (2019). Replication-coupled histone H3.1 deposition determines nucleosome composition and heterochromatin dynamics during Arabidopsis seedling development. The New phytologist 221, 385–398.

Bouveret, R., Schonrock, N., Gruissem, W., and Hennig, L. (2006). Regulation of flowering time by Arabidopsis MSI1. Development 133, 1693–1702.

Bowman, A., Ward, R., Wiechens, N., Singh, V., El-Mkami, H., Norman, D.G., and Owen-Hughes, T. (2011). The histone chaperones Nap1 and Vps75 bind histones H3 and H4 in a tetrameric conformation. Molecular cell 41, 398–408.

Bowman, A., Hammond, C.M., Stirling, A., Ward, R., Shang, W., El-Mkami, H., Robinson, D.A., Svergun, D.I., Norman, D.G., and Owen-Hughes, T. (2014). The histone chaperones Vps75 and Nap1 form ring-like, tetrameric structures in solution. Nucleic acids research 42, 6038–6051.

Citovsky, V., Lee, L.Y., Vyas, S., Glick, E., Chen, M.H., Vainstein, A., Gafni, Y., Gelvin, S.B., and Tzfira, T. (2006). Subcellular localization of interacting proteins by bimolecular fluorescence complementation in planta. Journal of molecular biology 362, 1120–1131.

D’Arcy, S., Martin, K.W., Panchenko, T., Chen, X., Bergeron, S., Stargell, L.A., Black, B.E., and Luger, K. (2013). Chaperone Nap1 shields histone surfaces used in a nucleosome and can put H2A-H2B in an unconventional tetrameric form. Molecular cell 51, 662–677.

Dellaporta, S.L., Wood, J., and Hicks, J.B. (1983). A plant DNA minipreparation: version II. Plant Mol. Biol. Report. 1, 19–21.

Dewari, P.S., and Bhargava, P. (2014). Genome-wide mapping of yeast histone chaperone anti-silencing function 1 reveals its role in condensin binding with chromatin. PloS one 9, e108652.

Dong, A., Liu, Z., Zhu, Y., Yu, F., Li, Z., Cao, K., and Shen, W.H. (2005). Interacting proteins and differences in nuclear transport reveal specific functions for the NAP1 family proteins in plants. Plant Physiol 138, 1446–1456.

Dorn, A., Feller, L., Castri, D., Rohrig, S., Enderle, J., Herrmann, N.J., Block-Schmidt, A., Trapp, O., Kohler, L., and Puchta, H. (2019). An Arabidopsis FANCJ helicase homologue is required for DNA crosslink repair and rDNA repeat stability. PLoS Genet 15, e1008174.

Dronamraju, R., Ramachandran, S., Jha, D.K., Adams, A.T., DiFiore, J.V., Parra, M.A., Dokholyan, N.V., and Strahl, B.D. (2017). Redundant Functions for Nap1 and Chz1 in H2A.Z Deposition. Scientific reports 7, 10791.

Duc, C., Benoit, M., Le Goff, S., Simon, L., Poulet, A., Cotterell, S., Tatout, C., and Probst, A.V. (2015). The histone chaperone complex HIR maintains nucleosome occupancy and counterbalances impaired histone deposition in CAF-1 complex mutants. The Plant journal: for cell and molecular biology 81, 707–722.

Duc, C., Benoit, M., Detourne, G., Simon, L., Poulet, A., Jung, M., Veluchamy, A., Latrasse, D., Le Goff, S., Cotterell, S., Tatout, C., Benhamed, M., and Probst, A.V. (2017). Arabidopsis ATRX Modulates H3.3 Occupancy and Fine-Tunes Gene Expression. The Plant cell 29, 1773–1793.

Dvorackova, M., Fojtova, M., and Fajkus, J. (2015). Chromatin dynamics of plant telomeres and ribosomal genes. The Plant journal: for cell and molecular biology 83, 18–37.

Eckert-Boulet, N., and Lisby, M. (2010). Regulation of homologous recombination at telomeres in budding yeast. FEBS letters 584, 3696–3702.

Endo, M., Ishikawa, Y., Osakabe, K., Nakayama, S., Kaya, H., Araki, T., Shibahara, K., Abe, K., Ichikawa, H., Valentine, L., Hohn, B., and Toki, S. (2006). Increased frequency of homologous recombination and T-DNA integration in Arabidopsis CAF-1 mutants. The EMBO journal 25, 5579–5590.

Exner, V., Taranto, P., Schonrock, N., Gruissem, W., and Hennig, L. (2006). Chromatin assembly factor CAF-1 is required for cellular differentiation during plant development. Development 133, 4163–4172.

Friesner, J.D., Liu, B., Culligan, K., and Britt, A.B. (2005). Ionizing radiation-dependent gamma-H2AX focus formation requires ataxia telangiectasia mutated and ataxia telangiectasia mutated and Rad3-related. Molecular biology of the cell 16, 2566–2576.

Fulcher, N., and Sablowski, R. (2009). Hypersensitivity to DNA damage in plant stem cell niches. Proceedings of the National Academy of Sciences of the United States of America 106, 20984–20988.

Fyodorov, D.V., and Kadonaga, J.T. (2003). Chromatin assembly in vitro with purified recombinant ACF and NAP-1. Methods in enzymology 371, 499–515.

Galbraith, D.W., Janda, J., and Lambert, G.M. (2011). Multiparametric analysis, sorting, and transcriptional profiling of plant protoplasts and nuclei according to cell type. Methods in molecular biology 699, 407–429.

Gao, J., Zhu, Y., Zhou, W., Molinier, J., Dong, A., and Shen, W.H. (2012). NAP1 family histone chaperones are required for somatic homologous recombination in Arabidopsis. The Plant cell 24, 1437–1447.

Gonzalez-Arzola, K., Diaz-Quintana, A., Rivero-Rodriguez, F., Velazquez-Campoy, A., De la Rosa, M.A., and Diaz-Moreno, I. (2017). Histone chaperone activity of Arabidopsis thaliana NRP1 is blocked by cytochrome c. Nucleic acids research 45, 2150–2165.

Hammond, C.M., Stromme, C.B., Huang, H., Patel, D.J., and Groth, A. (2017). Histone chaperone networks shaping chromatin function. Nature reviews. Molecular cell biology 18, 141–158.

Hisanaga, T., Ferjani, A., Horiguchi, G., Ishikawa, N., Fujikura, U., Kubo, M., Demura, T., Fukuda, H., Ishida, T., Sugimoto, K., and Tsukaya, H. (2013). The ATM-Dependent DNA Damage Response Acts as an Upstream Trigger for Compensation in the fas1 Mutation during Arabidopsis Leaf Development. Plant Physiol 162, 831–841.

Huang, T.H., Fowler, F., Chen, C.C., Shen, Z.J., Sleckman, B., and Tyler, J.K. (2018). The Histone Chaperones ASF1 and CAF-1 Promote MMS22L-TONSL-Mediated Rad51 Loading onto ssDNA during Homologous Recombination in Human Cells. Molecular cell 69, 879–892 e875.

Chen, X., D’Arcy, S., Radebaugh, C.A., Krzizike, D.D., Giebler, H.A., Huang, L., Nyborg, J.K., Luger, K., and Stargell, L.A. (2016). Histone Chaperone Nap1 Is a Major Regulator of Histone H2A-H2B Dynamics at the Inducible GAL Locus. Molecular and cellular biology 36, 1287–1296.

Chereji, R.V., Ocampo, J., and Clark, D.J. (2017). MNase-Sensitive Complexes in Yeast: Nucleosomes and Non-histone Barriers. Molecular cell 65, 565–577 e563.

Ishii, S., Koshiyama, A., Hamada, F.N., Nara, T.Y., Iwabata, K., Sakaguchi, K., and Namekawa, S.H. (2008). Interaction between Lim15/Dmc1 and the homologue of the large subunit of CAF-1: a molecular link between recombination and chromatin assembly during meiosis. The FEBS journal 275, 2032–2041.

Ivics, Z., and Izsvak, Z. (2010). Repetitive elements and genome instability. Seminars in cancer biology 20, 197–199.

Jacob, Y., Bergamin, E., Donoghue, M.T., Mongeon, V., LeBlanc, C., Voigt, P., Underwood, C.J., Brunzelle, J.S., Michaels, S.D., Reinberg, D., Couture, J.F., and Martienssen, R.A. (2014). Selective methylation of histone H3 variant H3.1 regulates heterochromatin replication. Science 343, 1249–1253.

Jin, C., Zang, C., Wei, G., Cui, K., Peng, W., Zhao, K., and Felsenfeld, G. (2009). H3.3/H2A.Z double variant-containing nucleosomes mark ‘nucleosome-free regions’ of active promoters and other regulatory regions. Nat Genet 41, 941–945.

Kaufman, P.D., Kobayashi, R., Kessler, N., and Stillman, B. (1995). The p150 and p60 subunits of chromatin assembly factor I: a molecular link between newly synthesized histones and DNA replication. Cell 81, 1105–1114.

Kaya, H., Shibahara, K.I., Taoka, K.I., Iwabuchi, M., Stillman, B., and Araki, T. (2001). FASCIATA genes for chromatin assembly factor-1 in arabidopsis maintain the cellular organization of apical meristems. Cell 104, 131–142.

Kirik, A., Pecinka, A., Wendeler, E., and Reiss, B. (2006). The chromatin assembly factor subunit FASCIATA1 is involved in homologous recombination in plants. The Plant cell 18, 2431–2442.

Kobayashi, T., Horiuchi, T., Tongaonkar, P., Vu, L., and Nomura, M. (2004). SIR2 regulates recombination between different rDNA repeats, but not recombination within individual rRNA genes in yeast. Cell 117, 441–453.

Kubik, S., Bruzzone, M.J., Albert, B., and Shore, D. (2017). A Reply to “MNase-Sensitive Complexes in Yeast: Nucleosomes and Non-histone Barriers,” by Chereji et al. Molecular cell 65, 578–580.

Kumar, A., Kumar Singh, A., Chandrakant Bobde, R., and Vasudevan, D. (2019). Structural Characterization of Arabidopsis thaliana NAP1-Related Protein 2 (AtNRP2) and Comparison with its Homolog AtNRP1. Molecules 24.

Lankenau, S., Barnickel, T., Marhold, J., Lyko, F., Mechler, B.M., and Lankenau, D.H. (2003). Knock-out targeting of the Drosophila Nap1 gene and examination of DNA repair tracts in the recombination products. Genetics 163, 611–623.

Levine, A.J., Ting, D.T., and Greenbaum, B.D. (2016). P53 and the defenses against genome instability caused by transposons and repetitive elements. BioEssays: news and reviews in molecular, cellular and developmental biology 38, 508–513.

Liu, X., Gao, L., Dinh, T.T., Shi, T., Li, D., Wang, R., Guo, L., Xiao, L., and Chen, X. (2014). DNA topoisomerase I affects polycomb group protein-mediated epigenetic regulation and plant development by altering nucleosome distribution in Arabidopsis. The Plant cell 26, 2803–2817.

Liu, Z., Zhu, Y., Gao, J., Yu, F., Dong, A., and Shen, W.H. (2009). Molecular and reverse genetic characterization of NUCLEOSOME ASSEMBLY PROTEIN1 (NAP1) genes unravels their function in transcription and nucleotide excision repair in Arabidopsis thaliana. The Plant journal: for cell and molecular biology 59, 27–38.

Luebben, W.R., Sharma, N., and Nyborg, J.K. (2010). Nucleosome eviction and activated transcription require p300 acetylation of histone H3 lysine 14. Proceedings of the National Academy of Sciences of the United States of America 107, 19254–19259.

Lundin, C., North, M., Erixon, K., Walters, K., Jenssen, D., Goldman, A.S., and Helleday, T. (2005). Methyl methanesulfonate (MMS) produces heat-labile DNA damage but no detectable in vivo DNA double-strand breaks. Nucleic acids research 33, 3799–3811.

Manova, V., and Gruszka, D. (2015). DNA damage and repair in plants - from models to crops. Frontiers in plant science 6, 885.

Mello, J.A., Sillje, H.H.W., Roche, D.M.J., Kirschner, D.B., Nigg, E.A., and Almouzni, G. (2002). Human Asf1 and CAF-1 interact and synergize in a repair-coupled nucleosome assembly pathway. Embo Reports 3, 329–334.

Moshkin, Y.M., Doyen, C.M., Kan, T.W., Chalkley, G.E., Sap, K., Bezstarosti, K., Demmers, J.A., Ozgur, Z., van Ijcken, W.F., and Verrijzer, C.P. (2013). Histone chaperone NAP1 mediates sister chromatid resolution by counteracting protein phosphatase 2A. PLoS Genet 9, e1003719.

Mozgova, I., Mokros, P., and Fajkus, J. (2010). Dysfunction of chromatin assembly factor 1 induces shortening of telomeres and loss of 45S rDNA in Arabidopsis thaliana. The Plant cell 22, 2768–2780.

Mozgova, I., Wildhaber, T., Trejo-Arellano, M.S., Fajkus, J., Roszak, P., Kohler, C., and Hennig, L. (2018). Transgenerational phenotype aggravation in CAF-1 mutants reveals parent-of-origin specific epigenetic inheritance. The New phytologist.

Mozgova, I., Wildhaber, T., Liu, Q., Abou-Mansour, E., L’Haridon, F., Metraux, J.P., Gruissem, W., Hofius, D., and Hennig, L. (2015). Chromatin assembly factor CAF-1 represses priming of plant defence response genes. Nature plants 1, 15127.

Muchova, V., Amiard, S., Mozgova, I., Dvorackova, M., Gallego, M.E., White, C., and Fajkus, J. (2015). Homology-dependent repair is involved in 45S rDNA loss in plant CAF-1 mutants. The Plant journal: for cell and molecular biology 81, 198–209.

Munoz-Viana, R., Wildhaber, T., Trejo-Arellano, M.S., Mozgova, I., and Hennig, L. (2017). Arabidopsis Chromatin Assembly Factor 1 is required for occupancy and position of a subset of nucleosomes. The Plant journal: for cell and molecular biology 92, 363–374.

Nakagawa, T., Bulger, M., Muramatsu, M., and Ito, T. (2001). Multistep chromatin assembly on supercoiled plasmid DNA by nucleosome assembly protein-1 and ATP-utilizing chromatin assembly and remodeling factor. The Journal of biological chemistry 276, 27384–27391.

Neplechova, K., Sykorova, E., and Fajkus, J. (2005). Comparison of different kinds of probes used for analysis of variant telomeric sequences. Biophysical chemistry 117, 225–231.

Otero, S., Desvoyes, B., Peiro, R., and Gutierrez, C. (2016). Histone H3 Dynamics Reveal Domains with Distinct Proliferation Potential in the Arabidopsis Root. The Plant cell 28, 1361–1371.

Ozer, O., and Hickson, I.D. (2018). Pathways for maintenance of telomeres and common fragile sites during DNA replication stress. Open biology 8.

Pavlistova, V., Dvorackova, M., Jez, M., Mozgova, I., Mokros, P., and Fajkus, J. (2016). Phenotypic reversion in fas mutants of Arabidopsis thaliana by reintroduction of FAS genes: variable recovery of telomeres with major spatial rearrangements and transcriptional reprogramming of 45S rDNA genes. The Plant journal: for cell and molecular biology 88, 411–424.

Pecinka, A., and Liu, C.H. (2014). Drugs for plant chromosome and chromatin research. Cytogenetic and genome research 143, 51–59.

Peterson, R., Slovin, J.P., and Chen, C. (2010). A simplified method for differential staining of aborted and non-aborted pollen grains. International Journal of Plant Biology 1, 13.

Pfab, A., Breindl, M., and Grasser, K.D. (2018). The Arabidopsis histone chaperone FACT is required for stress-induced expression of anthocyanin biosynthetic genes. Plant molecular biology 96, 367–374.

Pizzolato, J.F., and Saltz, L.B. (2003). The camptothecins. Lancet 361, 2235–2242.

Pontvianne, F., Blevins, T., Chandrasekhara, C., Mozgova, I., Hassel, C., Pontes, O.M., Tucker, S., Mokros, P., Muchova, V., Fajkus, J., and Pikaard, C.S. (2013). Subnuclear partitioning of rRNA genes between the nucleolus and nucleoplasm reflects alternative epiallelic states. Genes & development 27, 1545–1550.

Prasad, R., D’Arcy, S., Hada, A., Luger, K., and Bartholomew, B. (2016). Coordinated Action of Nap1 and RSC in Disassembly of Tandem Nucleosomes. Molecular and cellular biology 36, 2262–2271.

Ramirez-Parra, E., and Gutierrez, C. (2007). E2F regulates FASCIATA1, a chromatin assembly gene whose loss switches on the endocycle and activates gene expression by changing the epigenetic status. Plant Physiol 144, 105–120.

Rogner, U.C., Spyropoulos, D.D., Le Novere, N., Changeux, J.P., and Avner, P. (2000). Control of neurulation by the nucleosome assembly protein-1-like 2. Nature Genetics 25, 431–435.

Rohrig, S., Schropfer, S., Knoll, A., and Puchta, H. (2016). The RTR Complex Partner RMI2 and the DNA Helicase RTEL1 Are Both Independently Involved in Preserving the Stability of 45S rDNA Repeats in Arabidopsis thaliana. PLoS Genet 12, e1006394.

Roudier, F., Ahmed, I., Berard, C., Sarazin, A., Mary-Huard, T., Cortijo, S., Bouyer, D., Caillieux, E., Duvernois-Berthet, E., Al-Shikhley, L., Giraut, L., Despres, B., Drevensek, S., Barneche, F., Derozier, S., Brunaud, V., Aubourg, S., Schnittger, A., Bowler, C., Martin-Magniette, M.L., Robin, S., Caboche, M., and Colot, V. (2011). Integrative epigenomic mapping defines four main chromatin states in Arabidopsis. The EMBO journal 30, 1928–1938.

Sequeira-Mendes, J., Araguez, I., Peiro, R., Mendez-Giraldez, R., Zhang, X., Jacobsen, S.E., Bastolla, U., and Gutierrez, C. (2014). The Functional Topography of the Arabidopsis Genome Is Organized in a Reduced Number of Linear Motifs of Chromatin States. The Plant cell 26, 2351–2366.

Sfeir, A., Kosiyatrakul, S.T., Hockemeyer, D., MacRae, S.L., Karlseder, J., Schildkraut, C.L., and de Lange, T. (2009). Mammalian telomeres resemble fragile sites and require TRF1 for efficient replication. Cell 138, 90–103.

Shimada, M., Chen, W.Y., Nakadai, T., Onikubo, T., Guermah, M., Rhodes, D., and Roeder, R.G. (2019). Gene-Specific H1 Eviction through a Transcriptional Activator-->p300-->NAP1-->H1 Pathway. Molecular cell 74, 268–283 e265.

Schrumpfova, P.P., Fojtova, M., Mokros, P., Grasser, K.D., and Fajkus, J. (2011). Role of HMGB proteins in chromatin dynamics and telomere maintenance in Arabidopsis thaliana. Current protein & peptide science 12, 105–111.

Schrumpfova, P.P., Vychodilova, I., Dvorackova, M., Majerska, J., Dokladal, L., Schorova, S., and Fajkus, J. (2014). Telomere repeat binding proteins are functional components of Arabidopsis telomeres and interact with telomerase. The Plant journal: for cell and molecular biology 77, 770–781.

Schwabish, M.A., and Struhl, K. (2006). Asf1 mediates histone eviction and deposition during elongation by RNA polymerase II. Molecular cell 22, 415–422.

Slamenova, D., Dusinska, M., Bastlova, T., and Gabelova, A. (1990). Differences between survival, mutagenicity and DNA replication in MMS-and MNU-treated V79 hamster cells. Mutation research 228, 97–103.

Takeuchi, Y., Horiuchi, T., and Kobayashi, T. (2003). Transcription-dependent recombination and the role of fork collision in yeast rDNA. Genes & development 17, 1497–1506.

Tang, Y., Meeth, K., Jiang, E., Luo, C., and Marmorstein, R. (2008). Structure of Vps75 and implications for histone chaperone function. Proceedings of the National Academy of Sciences of the United States of America 105, 12206–12211.

Tyler, J.K., Collins, K.A., Prasad-Sinha, J., Amiott, E., Bulger, M., Harte, P.J., Kobayashi, R., and Kadonaga, J.T. (2001). Interaction between the Drosophila CAF-1 and ASF1 chromatin assembly factors. Molecular and cellular biology 21, 6574–6584.

Vannier, J.B., Depeiges, A., White, C., and Gallego, M.E. (2009). ERCC1/XPF protects short telomeres from homologous recombination in Arabidopsis thaliana. PLoS Genet 5, e1000380.

Varas, J., Santos, J.L., and Pradillo, M. (2017). The Absence of the Arabidopsis Chaperone Complex CAF-1 Produces Mitotic Chromosome Abnormalities and Changes in the Expression Profiles of Genes Involved in DNA Repair. Frontiers in plant science 8, 525.

Varas, J., Sanchez-Moran, E., Copenhaver, G.P., Santos, J.L., and Pradillo, M. (2015). Analysis of the Relationships between DNA Double-Strand Breaks, Synaptonemal Complex and Crossovers Using the Atfas1-4 Mutant. Plos Genetics 11.

Wollmann, H., Stroud, H., Yelagandula, R., Tarutani, Y., Jiang, D., Jing, L., Jamge, B., Takeuchi, H., Holec, S., Nie, X., Kakutani, T., Jacobsen, S.E., and Berger, F. (2017). The histone H3 variant H3.3 regulates gene body DNA methylation in Arabidopsis thaliana. Genome biology 18, 94.

Yelagandula, R., Stroud, H., Holec, S., Zhou, K., Feng, S., Zhong, X., Muthurajan, U.M., Nie, X., Kawashima, T., Groth, M., Luger, K., Jacobsen, S.E., and Berger, F. (2014). The histone variant H2A.W defines heterochromatin and promotes chromatin condensation in Arabidopsis. Cell 158, 98–109.

Zachova, D., Fojtova, M., Dvorackova, M., Mozgova, I., Lermontova, I., Peska, V., Schubert, I., Fajkus, J., and Sykorova, E. (2013). Structure-function relationships during transgenic telomerase expression in Arabidopsis. Physiologia plantarum 149, 114–126.

Zhang, Q., Giebler, H.A., Isaacson, M.K., and Nyborg, J.K. (2015). Eviction of linker histone H1 by NAP-family histone chaperones enhances activated transcription. Epigenetics & chromatin 8, 30.

Zhou, W., Gao, J., Ma, J., Cao, L., Zhang, C., Zhu, Y., Dong, A., and Shen, W.H. (2016). Distinct roles of the histone chaperones NAP1 and NRP and the chromatin-remodeling factor INO80 in somatic homologous recombination in Arabidopsis thaliana. The Plant journal: for cell and molecular biology 88, 397–410.

Zhu, Y., Dong, A., Meyer, D., Pichon, O., Renou, J.P., Cao, K., and Shen, W.H. (2006). Arabidopsis NRP1 and NRP2 encode histone chaperones and are required for maintaining postembryonic root growth. The Plant cell 18, 2879–2892.

Zhu, Y., Weng, M., Yang, Y., Zhang, C., Li, Z., Shen, W.H., and Dong, A. (2011). Arabidopsis homologues of the histone chaperone ASF1 are crucial for chromatin replication and cell proliferation in plant development. The Plant journal: for cell and molecular biology 66, 443–455.

